# Row1, a member of a new family of conserved fungal proteins involved in infection, is required for appressoria functionality in Ustilago maydis

**DOI:** 10.1101/2023.09.11.557153

**Authors:** María Dolores Pejenaute-Ochoa, Laura Tomás-Gallardo, José I. Ibeas, Ramón R. Barrales

## Abstract

The appressorium of phytopathogenic fungi is a specific structure with a crucial role in plant cuticle penetration. Pathogens with melanized appressoria break the cuticle through cell wall melanization and intracellular turgor pressure. However, in fungi with non-melanized appressorium, the mechanisms governing cuticle penetration are poorly understood. Here we characterize Row1, a previously uncharacterized appressoria-specific protein of *Ustilago maydis* that localizes to membrane and secretory vesicles. Deletion of *row1* decrease appressoria formation and plant penetration, thereby reducing virulence. Specifically, the Δ*row1* mutant has a thicker cell wall that is more resistant to glucanase degradation. We also observed that the Δ*row1* mutant has secretion defects. Our data suggest that Row1 could modify the glucans that form the fungal cell wall and may be involved in unconventional protein secretion, thereby promoting both appressoria maturation and penetration. We show that Row1 is functionally conserved at least among Ustilaginaceae and belongs to the Row family, which consists of five other proteins that are highly conserved among Basidiomycota fungi and are involved in *U. maydis* virulence. We observed similarities in localization between Row1 and Row2, which is also involved in cell wall remodelling and secretion, suggesting similar molecular functions for members of this protein family.

## INTRODUCTION

The interaction between plants and pathogenic fungi involves sophisticated mechanisms and elements from both organisms. Pathogens require invading strategies, such as the development of specialized structures to penetrate the plant cuticle (Ryder & Talbot, 2015; Shi *et al*., 2023) and a camouflage machinery to prevent recognition by their host (Uhse & Djamei, 2018; Yang, 2022). Plants prevent fungal infection by recognizing pathogen-associated molecular patterns (PAMPs) through their pattern recognition receptors (Boller & Felix, 2009), which activates PAMP-triggered immunity (Dodds & Rathjen, 2010; Uhse & Djamei, 2018). To counteract PAMP-triggered immunity, pathogens use effectors, secreted proteins that function either at the interface between host and pathogen or inside host cells (Lanver *et al*., 2017), which activate effector-triggered immunity to suppress host defences and support infection (Gupta *et al*., 2015).

This plant-pathogen interaction is highly dependent on the fungal cell wall, which is composed mainly of polysaccharides and proteins and serves as the initial barrier between the two organisms (Gow *et al*., 2017; Geoghegan *et al*., 2017; Tanaka & Kahmann, 2021). The inner layer of the cell wall consists predominantly of a chitin-glucan core (Latgé, 2007; Gow *et al*., 2017; Geoghegan *et al*., 2017). Glucan constitutes approximately 50–60% of the dry weight of fungal cell wall and consists mainly of long linear chains of beta-1,3-linked glucose (Bowman & Free, 2006; Garcia-Rubio *et al*., 2019), while chitin is present in lower abundance (10–20%) as beta-1,4-linked chains of N-acetylglucosamine (Latgé, 2007; Gow *et al*., 2017; Garcia-Rubio *et al*., 2019). The outer layer includes mannosylated proteins representing 20-30% of fungal cell wall (Bowman & Free, 2006), and most of them are anchored to the plasma membrane by the lipid glycosylphosphatidylinositol (GPI) (De Groot *et al*., 2005; Vogt *et al*., 2020). Recent studies suggest that the GPI anchoring of these proteins is crucial for their function, since loss of GPI compromises cell wall integrity and virulence in fungal pathogens (Rittenour & Harris, 2013; Liu *et al*., 2020). Moreover, the sugar fraction of the mannosylated proteins is also essential for virulence, since alterations of their N-or O-glycosylation pattern suppresses plant antifungal protein binding and killing activity, and plant infection (Fernández-Álvarez *et al*., 2009; 2012; 2013; Pejenaute-Ochoa *et al*., 2021; Ma *et al*., 2023). Given the location of mannoproteins on the surface of the cell wall, they are thought to be involved in host adhesion, evasion of the host immune response, and maintenance of cell wall integrity (Bowman & Free, 2006; Gow *et al*., 2017; Garcia-Rubio *et al*., 2019).

During the first stages of pathogenesis, fungi undergo dynamic cell wall remodelling, which is facilitated by the activity of glycohydrolases (such as chitinases and glucanases), chitin-deacetylases and transglycosylases (Gow *et al*., 2017; Geoghegan *et al*., 2017; Gow & Lenardon, 2023). The absence of many of these enzymes drastically reduces virulence (Mouyna *et al*., 2005; Wawra *et al*., 2016; Samalova *et al*., 2017; Bi *et al*., 2021). This remodelling mechanism enables the fungus to evade plant defence molecules (van den Burg *et al*., 2006; Mentlak *et al*., 2012; Geoghegan *et al*., 2017) and undergoes morphological changes to develop filaments, penetrate the plant cuticle, invade host tissues and colonize successfully (Mendgen *et al*., 1996; Wang & Lin, 2012; Lin *et al*., 2014). An essential morphological transition for establishing virulence in pathogenic fungi is the formation of the appressorium, a specialized structure that facilitates breaching of the plant cuticle, allowing effective colonization of the host (Ryder & Talbot, 2015; Chethana *et al*., 2021; Ryder *et al*., 2022).

Different types of appressoria and their mechanisms of penetration have been studied extensively in plant pathogens (O’Connell & Panstruga, 2006; Ryder & Talbot, 2015; Talbot, 2019; Chethana *et al*., 2021). Dark appressoria in fungi like *Colletotrichum* or *Magnaporthe* species are crucial for the infection process (de Jong *et al*., 1997; Perfect *et al*., 1999; Tucker & Talbot, 2001) and require cell wall melanization and glycerol accumulation for penetration (Mendgen *et al*., 1996; de Jong *et al*., 1997; Wilson & Talbot, 2009). Appressoria melanization and turgor pressure, which is coordinated with the secretion of plant-cell wall degrading enzymes (PCWDEs) (Presti *et al*., 2015; Wang & Wang, 2018; Yang, 2022), enable the fungus to mechanically breach the host surface. Other cereal pathogens, such as *Blumeria graminis, Phakopsora pachyrhizi* and *Ustilago maydis*, have non-melanized or slightly melanized appressoria (Mendgen *et al*., 1996; Lanver *et al*., 2014; Chethana *et al*., 2021; Ryder *et al*., 2022). In these pathogens, secretion of PCWDEs and effectors is particularly important for host invasion and for establishing disease progression (Kubicek *et al*., 2014; Bradley *et al*., 2022).

In the plant pathogenic fungus *U. maydis*, many effectors and PCWDEs have been identified and characterized (for examples see Lanver et al., 2017; Zuo et al., 2019; Ludwig et al., 2021; Navarrete et al., 2021; Moreno-Sánchez et al., 2021; Ökmen et al., 2022; Bindics et al., 2022). When this fungus penetrates the plant, it interacts closely with the surrounding plant plasma (Doehlemann et al., 2008), where it secretes proteins that modify host cell structure and function (Win *et al*., 2012; Lanver *et al*., 2017). The fungus also modifies its own cell wall to improve its infectivity and to evade the plant immune system (Mueller *et al*., 2008; Ruiz-Herrera *et al*., 2008; Lanver *et al*., 2014). To do this, *U. maydis* uses several strategies, such as converting chitin to chitosan (Rizzi *et al*., 2021; Ma *et al*., 2023), redecorating the surface of the hyphae by blocking plant antifungal activity (Ma *et al*., 2018), and reorganizing the fungal cell wall structure (Tanaka *et al*., 2020).

In this study, we characterize the protein Row1, remodelling of fungal cell wall 1. Here we show that deletion of *row1* leads to defects in appressoria formation and cell wall structure and disrupts normal protein secretion. Moreover, we demonstrated that Row1 belongs to a conserved protein family of five members that are also involved in pathogenesis.

## MATERIALS AND METHODS

### Plasmids and strain constructions, growth conditions and infection assays

All *U. maydis* strains used in this study are listed in Supporting Information Table S1. Southern Blot analysis was used to verify all deletion and complementation mutants as previously described (Moreno-Sánchez *et al*., 2021). Primers and plasmids used in this study, and the cloning procedure used to generate them, are listed in Table S2. Detailed cloning and strain generation, growth conditions and virulence assay are provided in **Methods S1**. Gene accession number is provided in Table S3.

### Adhesion and stress assays

Cell stress and cell wall integrity assays were developed with cultures grown at 28°C to exponential phase in CMD-2%glucose and spotted at OD_600_ of 0.4 onto CM plates supplemented with different stress-inducing agents. Specifically, Tunicamycin 1 μg/ml (Sigma-Aldrich) were used for reticulum stress, calcofluor white 10 μg/ml (Sigma-Aldrich) and Congo Red 10mM (Sigma-Aldrich) for cell wall integrity, Sorbitol 1M (Sigma-Aldrich, and NaCl 0.5M (Sigma-Aldrich) for osmotic pressure, and H_2_O*_2_* 0.75 mM (Sigma-Aldrich) for oxidative stress. Plates were incubated for 48 h at 28°C. Adhesion assay was performed as previously described (Fernández-Álvarez *et al*., 2012).

### Fungal Biomass Analysis

For fungal biomass quantification, 2cm long segments from the 3^rd^ leaf of 8 different plants at 2, 4 and 6 dpi were cut 1cm below the infective puncture and treated as previously described (Marín-Menguiano *et al*., 2019). 60 ng of total DNA was used as template for each reaction.

### Samples preparation for microscopy analysis

For hyphae proliferation, infected leaves from 1, 3 and 5 dpi were stained with wheat germ agglutinin-propidium iodide WGA-PI. Infected plants were distained with ethanol, treated 4h at 60°C with 10% KOH, washed in PBS1X buffer and then stained with PI to visualize plant tissues in red and WGA-AF488 to visualize the fungus in green.

To detect chitin, filaments induced for 5 hours were stained with WGA as explained (Fernández-Álvarez *et al*., 2009). To visualize filamentation and septa formation, cells were centrifugated and resuspended in Calcofluor White (CFW) staining solution (4 µg/mL CFW). For appressoria formation, infected leaves were stained with CFW (0.1 mg/mL) and observed 18h after plant inoculation. Chlorazole Black staining was performed as described (Brachmann et al., 2003) in leaves collected at 1dpi. All samples were observed using Delta Vision microscopy.

For Lallzyme treatment, filament cultures induced for 5 hours were resuspended in cold Lallzyme MMX® (0.015g/ml) as indicated (Fernández-Álvarez *et al*., 2013). Samples from all the examined strains were collected at 15 minutes and subjected to microscopic imaging using a DeltaVision microscope.

For transmission electron microscopy, samples were fixed, processed, and examined using a Zeiss Model Libra 120 transmission electron microscope in the General Research Services of the University of Seville (CITIUS).

Row family proteins tagged with GFP or mCherry were observed using DeltaVision microscope.

The features, filters and settings for microscopy are detailed in **Methods S1.**

### Protein and blotting assays

For protein secretion extraction, proteins in supernatants were collected after Trichloroacetic (TCA) – deoxycholate (DOC) precipitation. For cytosolic protein extraction, pellets were ground into a powder using a mortar/pestle under liquid nitrogen and were resuspended in lysis buffer (20 mM Tris-HCl, 0.5 M NaCl, pH 7.4) with protease inhibitor cocktail and centrifuged at 14000 rpm for 30 min at 4°C and supernatant was collected and quantified. 60 μg of each protein fraction was separated by SDS-PAGE and detected by western blot analysis.

Changes in protein secretion were relative quantified with the isobaric standard tandem tag (TMT) 10 plex labelling kit (Thermo Fisher Scientific).

Colony secretion assay was performed as previously described (Moreno-Sánchez *et al*., 2021).

The detailed protocols of protein extraction and Mass Spectometry assay, western blotting and data analysis can be found in **Methods S1.**

### Sequence Alignment, Phylogenetic Analysis and Predictive analysis tool

BlastP was used to search for Row1 homologues sequences in *U. maydis* and other fungi. For the Row1 Ustilaginaceae phylogenetic tree, reciprocal best hits blast was used. Multiple sequence alignments were generated by MAFFT v7 and visualized using Jalview. Phylogenetic analysis is explained in **Methods S1.** Predictive analysis tool used to infer proteins characteristic is thoroughly explained in **Methods S1**.

## RESULTS

### Row1 plays a role in appressoria progression inside plant tissues

We previously identified several *U. maydis* glycoproteins with effects on plant infection (Marín-Menguiano *et al*., 2019). Umag_00309, hereafter Row1 (remodelling of fungal cell wall protein 1), was an uncharacterized protein with no clear homology with previously characterized proteins. To confirm the relevance of Row1 in pathogenesis, we infected maize plants with two independent clones of *row1* deletion mutants in the sexually compatible *U. maydis* strains FB1 (a1b1) and FB2 (a2b2) (Banuett & Herskowitz, 1989)(Fig. S1a). We also performed Δ*row1* infection assays in the solopathogenic *U. maydis* strain SG200 (Fig. S1b) (Bölker *et al*., 1995), and in the CL13 strain (Fig. S1c), a progenitor strain of SG200 that has attenuated virulence (Bölker *et al*., 1995). Deletion of *row1* compromises infection in all these strains. As the results showed a greater effect in the CL13 background (Fig. S1), we reintroduced the *row1* allele in the CL13 Δ*row1* mutant and observed a full recovery of its virulence capacity, confirming a role for Row1 in infection (Fig. 1a). To ascertain the role of Row1 in pathogenesis, we first evaluated if loss of *row1* leads to growth defects under axenic conditions. We found no differences between the wild-type (WT) and a Δ*row1* mutant strain in generation time (Fig. S2a), cell morphology and length (Fig. S2b), or cellular adhesion ability (Fig. S2c). Furthermore, the Δ*row1* and WT strains responded similarly to oxidative, saline and cell wall stresses (Fig. S3). These results indicate that pathogenic defects in Δ*row1* infections are unlikely to be associated with defects in non-pathogenic cell cycle progression, which suggests that Row1 may be essential specifically to virulence. In agreement with this idea, we observed that *row1* is induced during infection at 1 day post-inoculation (dpi) (Fig. 1b), which is consistent with the previously developed global genomic profile of *U. maydis* (Lanver *et al*., 2018).

**Fig. 1.**
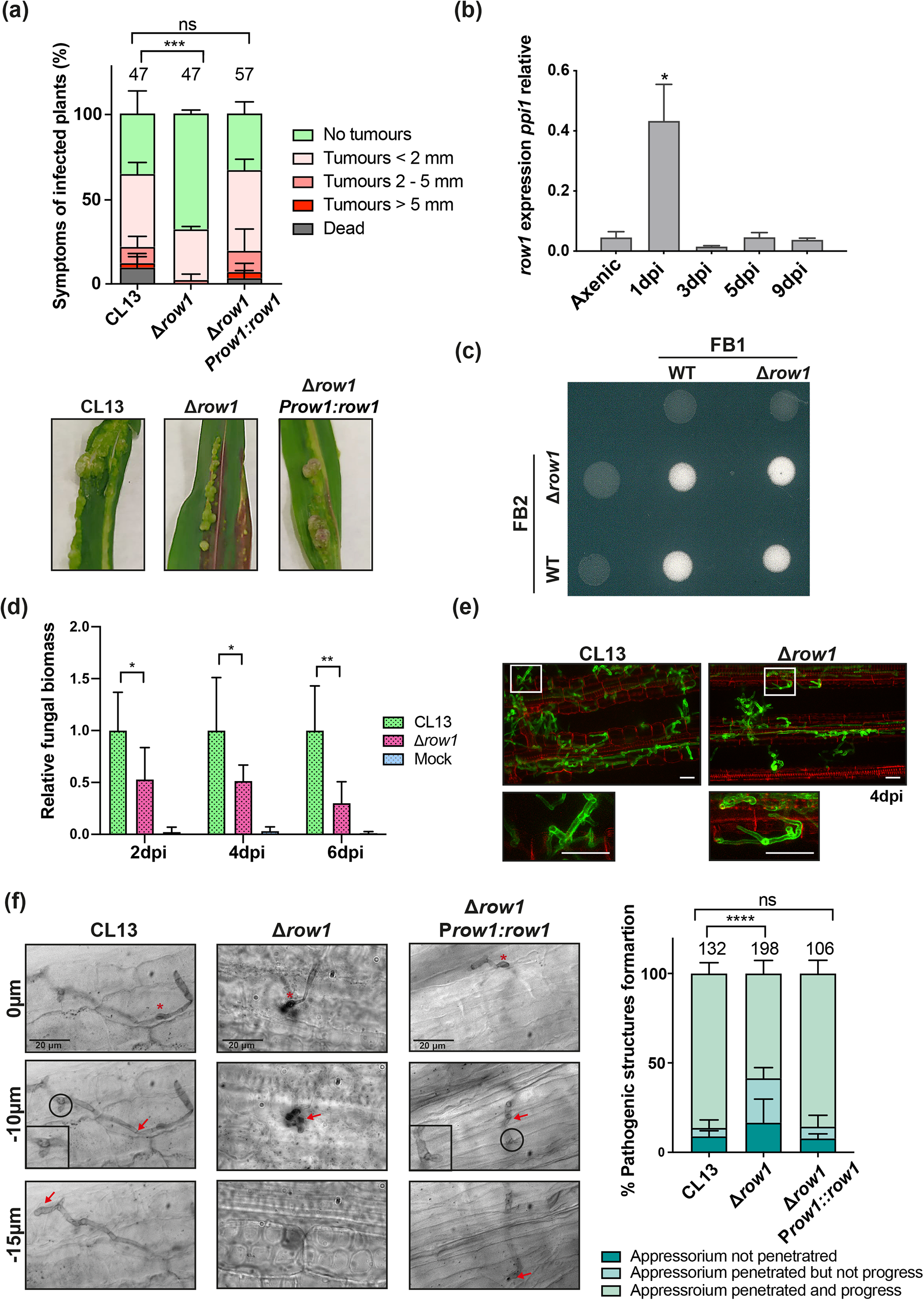
Row1 is important for appressoria progression by facilitating successful host tissue colonization and subsequent tumour formation. (a) Quantification of symptoms for plants infected with indicated strains at 14 dpi (left panel). Total number of infected plants is indicated above each column. Error bars represent the standard deviation from three independent replicates. The Mann–Whitney statistical test was performed for each mutant versus the WT strain (ns, not significant; \*\*\**p*-value < 0.005). Representative images of the most prevalent tumour category are shown in the right panel. (b) row1 expression levels relative to those of the *ppi1* gene measured by qRT-PCR. Error bars represent the standard deviation from three independent replicates. Student’s t-test statistical analysis was performed (\**p*-value < 0.05). **(c)** Mating assay of the compatible *U. maydis* strains (FB1, FB2, FB1 Δ*row1* and FB2 Δ*row1*) spotted alone or in combination and incubated on PD-charcoal plates. The white fuzzy appearance of the filaments is indicative of successful mating and dikaryotic hyphae formation **(d)** Fungal relative biomass was calculated by comparing the *U. maydis ppi1* gene and the *Z. mays gadph* gene and was measured using RT-qPCR of genomic DNA extracted from leaves infected with WT, Δ*row1* mutant or water as mock treatment at 2, 4, and 6 dpi. Error bars represent the standard deviation from four independent replicates. Student’s t-test statistical analysis was performed (**p-value < 0.005; \**p*-value < 0.05). **(e)** Maize leaves from plants infected with WT and the Δ*row1* mutant at 4 dpi were stained with propidium iodide (red) for plant cell visualization and with WGA-AF-488 (green) for *U. maydis* hyphae visualization by fluorescence microscopy. Scale bar represents 20 μm. **(f)** Leaf samples infected with WT and the Δ*row1* mutant were stained with Chlorazol Black and analysed by light microscopy 29 h after infection. In the left panel, the z-axis image projections show the site of appressorium formation and penetration (red asterisk), the hyphae invading the plant cells (red arrows), and the clamp cells (black circle). In the right panel, the identified structures are quantified in each strain. Error bars represent the standard deviation from three independent replicates. The total number of infected plants is indicated above each column. The Mann–Whitney statistical test was performed for each versus the WT strain (ns, not significant; \*\*\*\**p*-value < 0.0001).

As the first stages of the *U. maydis* pathogenic program require FB1xFB2 mating, we evaluated mating and filament capacity. However, we found no significant differences between WT and mutant strains (Fig. 1c). Because Row1 is not required for mating and its role in infection is more relevant in the CL13 background (Fig. 1a and Fig. S1c), we used this strain, which facilitates the detection of modest differences in virulence (Di Stasio *et al*., 2009; Djamei *et al*., 2011), for further infection experiments. First, we studied the mutant’s ability to proliferate inside the plant by analysing fungal biomass at 2, 4, and 6 dpi. We observed at least 50% less fungal biomass in the mutant compared to WT at all tested points (Fig. 1d). As we did not observe any structural defects in the proliferative hyphae of the Δ*row1* mutant (Fig. 1e), we evaluated whether its reduced abundance inside the host might be attributed to problems occurring at an earlier step in its pathogenic development. Therefore, we studied filamentation, appressoria formation, hyphal branching, and clamp cell formation by staining the fungus with Chlorazol Black at 29 hours post-infection. Although we detected no major morphological differences in these structures between the mutant and WT strains, approximately 42% of the appressoria of the Δ*row1* mutant did not penetrate or were arrested after penetration, in contrast to the 15% of those of the WT strain (Fig. 1f). Our findings suggest that Row1 is important for appressoria progression, which facilitates successful host tissue colonization and subsequent tumour formation.

### Row1 localizes to the secretory membrane system and accumulates at the appressorium during the initial stages of host interaction

We next aimed to uncover the role of Row1 in these pathogenesis defects. Using different databases, we identified Row1 as a GPI effector protein comprising a signal peptide (amino acids 1–21), a serine-rich region (297–401) with at least 14 putative mannosylation sites, and an alpha-helix transmembrane domain (403–423) that exposes the C-terminal domain of the protein to the extracellular region. Although we could not predict well-defined structures or the signal peptide, the Ser-rich domain or the transmembrane region, we predicted a globular structure with a central β-sheet in the central domain (amino acids 100–300) (Fig. 2a). As we did not identify any functional domains in Row1, we used the Sma3 tool (Casimiro-Soriguer *et al*., 2017), based on high-throughput annotation, to determine the protein’s potential function, cellular localization, biological process or protein structure. GO term annotation identified a putative role for Row1 in the polysaccharide catabolic process (GO:0000272), the xylan catabolic process (GO:0045493), hydrolase activity on glycosyl bonds (GO:0016798) and transmembrane transport (GO:0055085). To complement this information, we also studied protein localization. As previous data showed that *row1* is expressed at the beginning of the pathogenic program, we expressed *Row1* labelled with green fluorescent protein (GFP) under its own promoter in the AB33 strain, which harbours the compatible bE2/bW1 genes under the control of the nitrate-inducible *nar1* promoter.

**Fig. 2.**
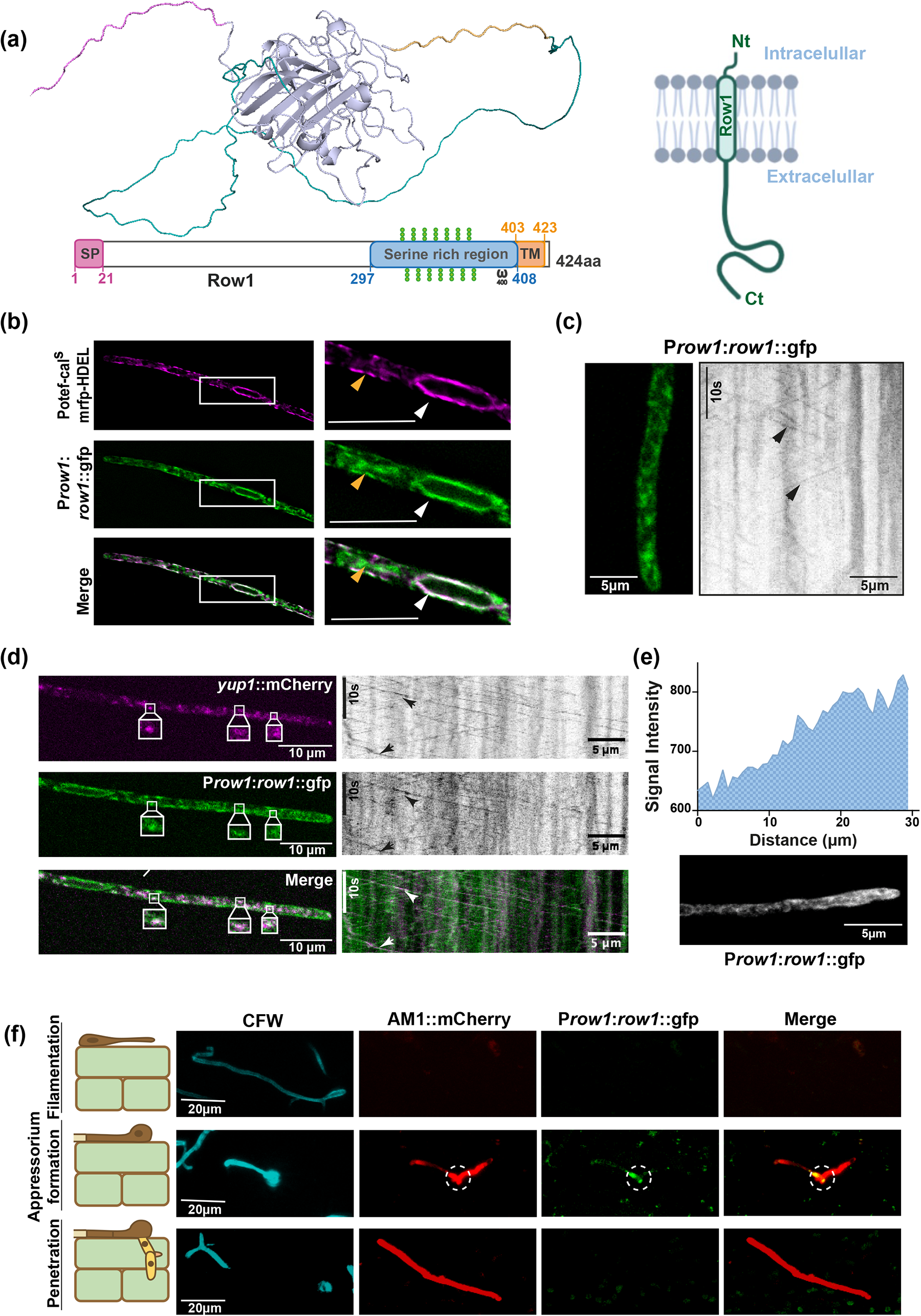
Row1 localizes in the secretory pathways and is accumulated at the appressorium. **(a)** 3D structure and domains of Row1. The 3D structure was obtained using AlphaFold. Left panel: Row1 has a signal peptide (SP; amino acids 1–21), a Ser-rich domain (amino acids 297–408) with several O-glycosylation sites (marked with three green spheres), a ω GPI anchor site (amino acid 400) and a transmembrane region (TM; amino acids 403–423). Right panel: Row1 is anchored to the plasma membrane through its TM region. The C-terminal region faces the extracellular space, while the N-terminal region faces the intracellular space. **(b)** Row1::GFP co-localization with the ER marker mRFP:HDEL is indicated by white arrows. Additional vesicles are marked with orange arrows. Scale bar represents 10 μm. **(c)** Row1 localization on hypha (left panel) and Row1 vesicle movement across time indicated by black arrows (right panel). **(d)** Row1::GFP partial co-localization with Yup1::mCherry vesicles (left panel). The kymographs show vesicle movement over time (right panel). **(e**) Quantification of Row1::GFP signal intensity (upper panel) along the growing hypha (lower panel). **(f)** Co-localization of Row1 with the appressorium using the AM1-mCherry reporter, which is specifically expressed in the tips of filaments that differentiate to an appressorium. Infected leaves were stained with CFW and observed by confocal microscopy 18 h after infection. The appressorium is distinguished by the formation of a characteristic red crook-like structure.

In this strain, filamentation, one of the first steps of the pathogenic program, is induced in nitrate-containing medium (Brachmann *et al*., 2001). When filamentation was induced, Row1 localized at the endoplasmic reticulum (ER) and plasma membrane, co-localizing with the ER marker mRFP::HDEL (Fig. 2b). In addition, Row1 was detected as small dots with bidirectional movement along defined cellular tracks, reminiscent of secretory vesicles (Fig. 2c). We confirmed the localization of Row1 in secretory vesicles by colocalization with Yup1 (Fig. 3d), a protein receptor (t-SNARE) involved in membrane fusion that is necessary for the delivery of cell wall components (Wedlich-Söldner *et al*., 2000; Fuchs *et al*., 2006). We also observed that Row1 accumulates at sites of active growth (Fig. 3e). Considering that *row1* is induced during the early stages of infection, alongside appressoria formation, and that Δ*row1* cells exhibit defects in appressoria formation, we hypothesized that the primary function of Row1 might be during appressorium formation. Thus, we examined Row1::GFP localization during appressorium formation by co-localization with the AM1::mCherry reporter, which is specifically expressed in the tips of filaments that are differentiating to appressoria. Our findings revealed specific Row1::GFP localization at appressoria, with no signal in the filament before or after appressorium formation (Fig. 2f).

**Fig. 3.**
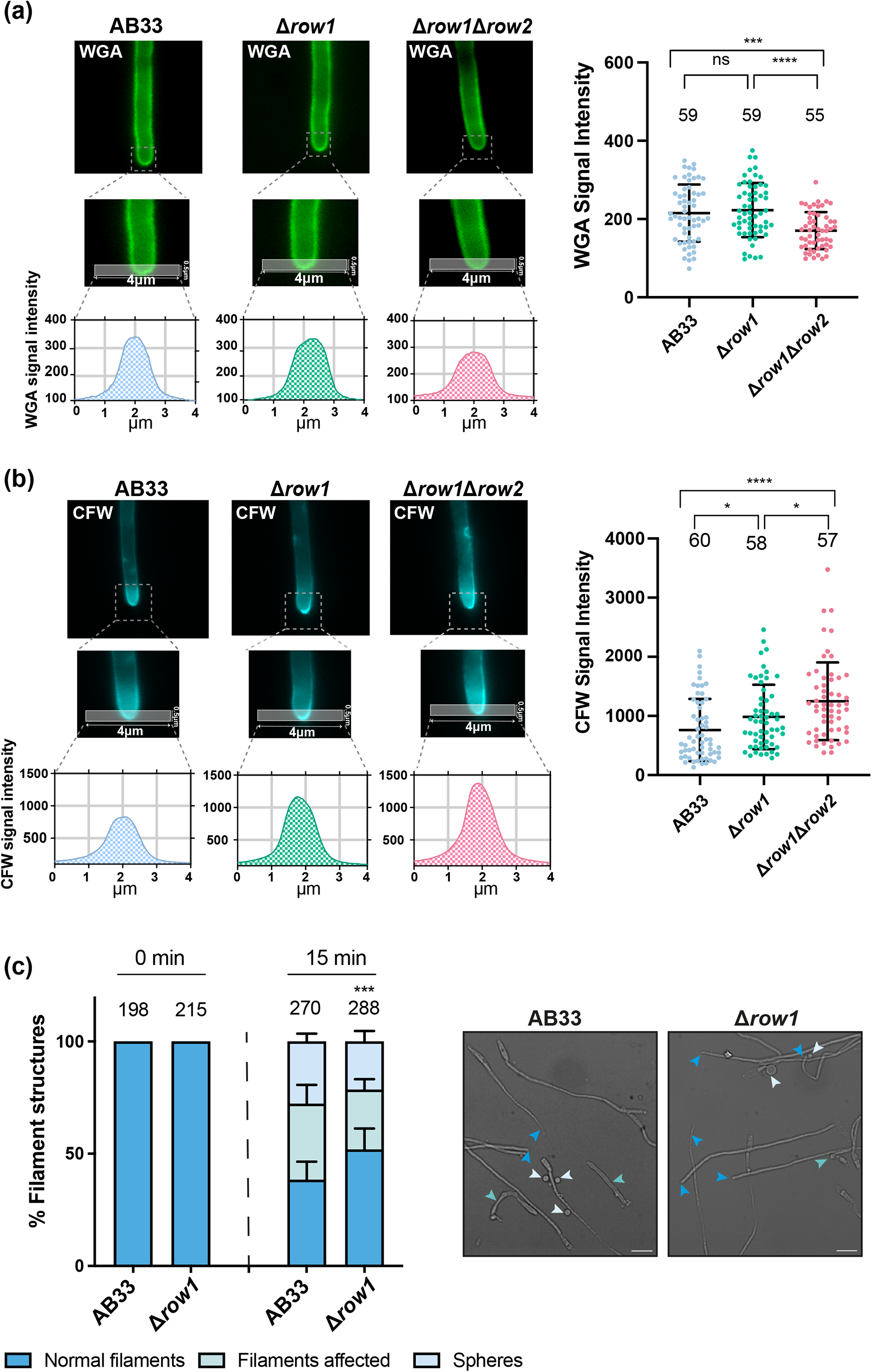
Row1 is essential for proper cell wall architecture. **(a)** Representative images of WGA-AF488 chitin staining in AB33, Δ*row1* and Δ*row1*Δ*row2* hyphae after 5 h of induction visualized by confocal microscopy. WGA signal intensity was quantified using the FIJI software, drawing a line at the hypha tip with the width and length indicated in the figure. This analysis was performed on a single z-plane from each image by calculating the average value for each point collected along the line and represented in the graphs (left panel). The graph in the right panel represents the maximum intensity for each filament, obtained by subtracting the background intensity. Scale bar represents 10 μm. The total number of filaments is indicated above each column. Error bars represent the standard deviation from three independent replicates. The Student’s t-test statistical analysis was performed (ns, no significant; \*\*\**p*-value < 0.0005; \*\*\**p*-value < 0.0001). **(b)** Representative images of CFW glucan staining in AB33, Δ*row1* and Δ*row1*Δ*row2* hyphae after 5 h of induction visualized by confocal microscopy. The quantification and analysis of the CFW signal followed the same procedure as in panel (a). Scale bar represents 10 μm. The total number of filaments is indicated above each column. Error bars represent the standard deviation from three independent replicates. The Student’s t-test statistical analysis was performed (ns not significant; \**p*-value < 0.05; \*\*\*\**p*-value < 0.0001).

### Row1 is essential for proper cell wall architecture

Based on the localization data and our prediction that Row1 is an effector protein with a potential role in polysaccharide degradation, we postulated that Row1 may function as a secreted PCWDE involved in facilitating successful penetration. To explore this possibility, we used a colony secretion assay (Krombach *et al*., 2018) in an SG200 background and induced the virulence program by growing the cells on Potato Dextrose (PD)-Charcoal media. However, no Row1::GFP signal was detected in either the pathogenic or non-pathogenic conditions (Fig. S4a). In addition, in a western blot assay, we observed the signal for Row1::GFP in the cytosolic lysate under induction conditions (Fig. S4b) but not in the secreted fraction. However, we detected several bands that might indicate the secretion and processing of Row1 (Fig. S4b). As we could not conclusively determine that Row1 is secreted, and many GPI proteins, as Row1 is predicted to be, are involved in fungal cell wall remodelling (Mouyna *et al*., 2005; Samalova *et al*., 2017; Bi *et al*., 2021), we explored its role in fungal cell wall remodelling.

To study cell wall composition, we stained filaments with the lectin WGA, which specifically binds to the N-acetylglucosamine monomers that form chitin (Nagata & Burger, 1974), conjugated to Alexa Fluor 488 (WGA-AF488). As has been previously observed (Flor-Parra *et al*., 2007), the WT filaments accumulated chitin at the growing hyphal tip, which was restricted to the growing apex. However, WT and mutant strains showed similar levels of accumulation (Fig. 3a). To determine if any other component of the cell wall was affected, we stained the hyphae with calcofluor white (CFW). CFW has affinity for the β-(1,4) glucans that connect the N-acetylglucosamine monomers, rather than the monomers themselves. It also demonstrates affinity for β-(1,3) glucans (Rasconi et al., 2009) . In this case, we found a stronger CFW signal in the Δ*row1* mutant than in the WT (Fig. 3b). At the same time, Δ*row1* hyphae were more resistant than WT hyphae to Lallzyme MMX, a mix of glucan digestion enzymes (Fig. 3c). These data could suggest that the loss of *row1* mainly affects glucan composition or the structure of the cell wall during hyphae growth. We next aimed to characterize the alterations in the cell wall that arise from *row1* deletion using transmission electron microscopy. Although the glycoprotein-rich outer layer of the cell wall showed no notable difference between the WT and mutant strains, the glucan-chitin inner layer showed a brighter signal and was thicker in the Δ*row1* mutant (Fig. 4), indicating a different cell wall structure.

**Fig. 4.**
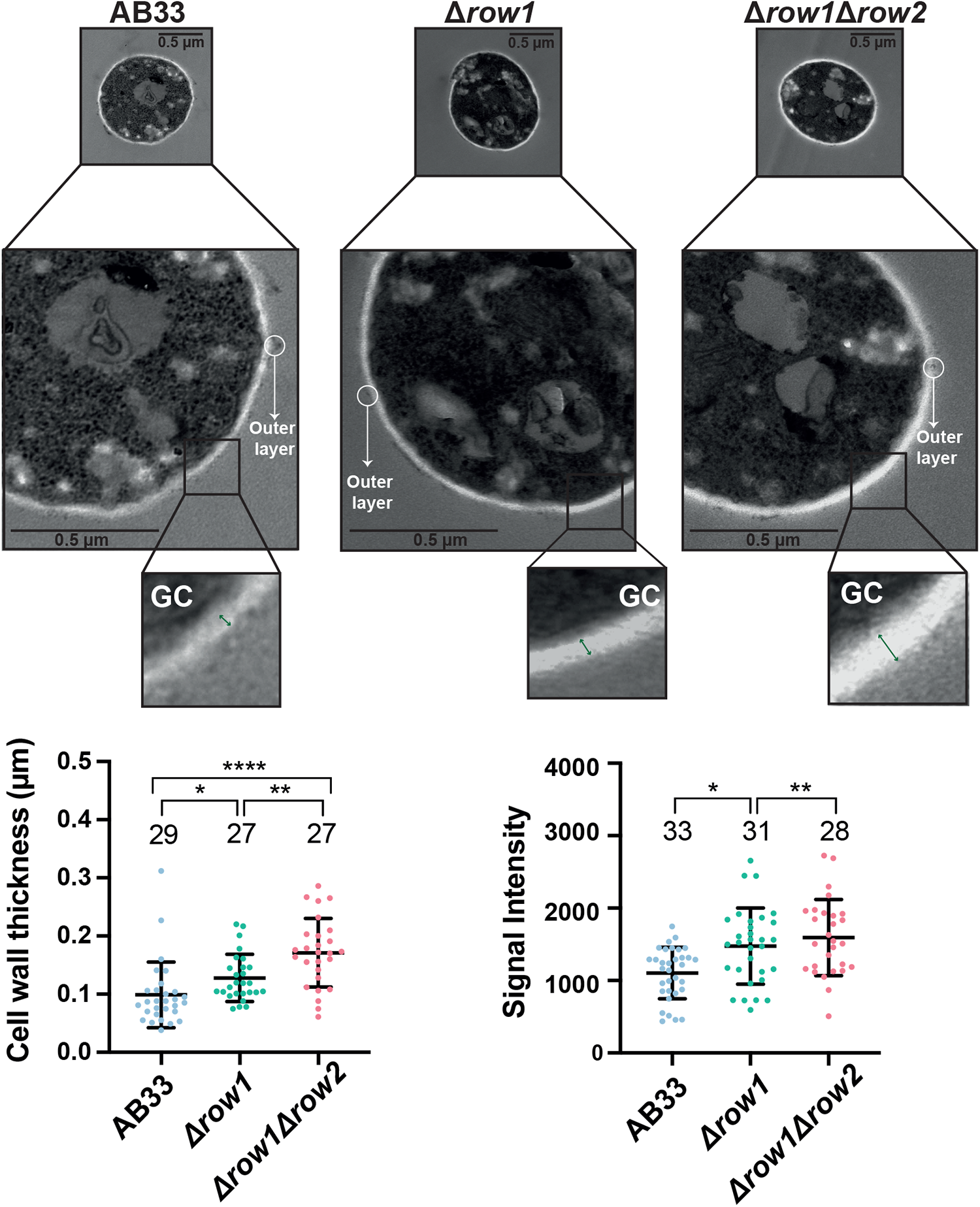
Δrow1 has a brighter and thicker inner layer and greater resistance to glucan degradation. **(a)** Upper panel: Transmission electron microscopy examination of AB33, Δ*row1* and Δ*row1*Δ*row2* cell walls from filament cultures grown for 5 h. The outer layer contains mannoproteins (white circles), while the inner layer contains glucan and chitin (GC). Lower panel: The graphs represent the measurements of the inner layer’s width and brightness intensity. The total number of filaments is indicated above each column. Error bars represent the standard deviation from three independent replicates. The Student’s t-test statistical analysis was performed for each mutant (ns, not significant; \**p*-value < 0.05; \*\**p*-value < 0.005; \*\*\*\**p-value <* 0.0001). **(b)** Left panel: Quantification of filament structures formed during a time course of protoplast formation. Filaments were treated with Lallzyme at room temperature. Protoplast formation was observed by wide-field microscopy, and the indicated structures were quantified (left panel). The total number of filaments is indicated above each column. Error bars represent the standard deviation from three independent replicates. Scale bar represents 10 μm. The Student’s t-test statistical analysis was performed (ns; no significant; ***p-value < 0.0005). Right panel: Representative images of the different filamentous structures formed at 15 min: normal filaments (blue arrows), filaments affected by the treatment (green arrows), and protoplasts or spheres (white arrows).

Next, we assessed whether the changes in cell wall composition and structure affected filament length and morphology. We measured filament length and counted bipolar or irregularly shaped hyphae in WT and the Δ*row1* mutant. The filaments in both strains had similar length and morphology (Fig. S5). These findings suggest that although the proper length and morphology of the pathogenic filaments do not require Row1, the loss of this protein leads to alterations in normal cell wall structure during the induction of the virulence program.

### Row1 is involved in faithful appressorium formation and maintenance of its cell-wall characteristics

Since appressorium formation involves a substantial transformation of the fungal cell wall, which transitions from a hyphal morphology to a dome-like structure (Ryder & Talbot, 2015), we hypothesized that alteration of fungal cell wall features might compromise the proper formation of this specialized structure. To investigate appressorium formation *in vivo* upon the loss of *row1*, we used the SG200 strain, cells of which form easily observable appressoria without mating. We quantified cells, filaments without appressorium, and filaments with appressorium in the WT and Δ*row1* mutant strains. While no differences in filamentation between the strains were detected (Fig. 5a, left), we found a lower percentage of filaments forming appressoria in the Δ*row1* mutant strain than in the WT strain (Fig. 5a, right). In agreement with our previous results, the appressoria exhibited a higher CFW signal in the Δ*row1* strain than in the WT strain (Fig. 5b). Our findings indicate that Row1 is important for the appressorium, potentially by remodelling the appressorium wall, which necessary for its formation and progression inside plant tissues.

**Fig. 5.**
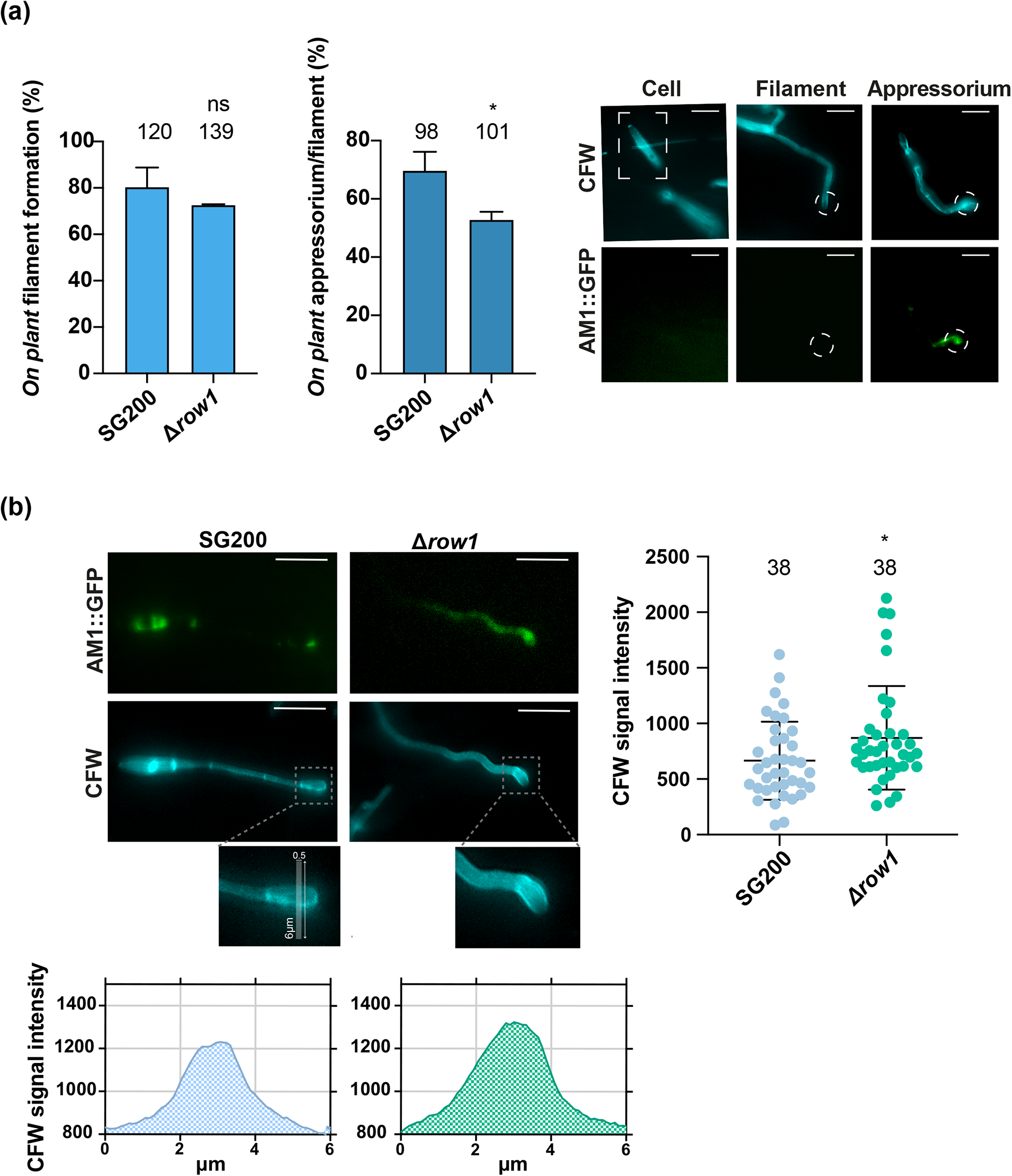
Row1 plays a role in appressoria formation and maintenance of its cell-wall characteristics. (**a**) Frequency of filaments and appressoria formation on plants using strains carrying the appressorium-specific AM1::GFP marker. Plants were infected with the indicated strains, stained after 18 h with calcofluor white (CFW) and then analysed for AM1 marker expression (AM1::GFP). The graph shows the results of two independent experiments, and the total number of structures is indicated above each column. The Mann–Whitney statistical test was performed for the mutant versus WT strain (ns, no significant; \**p*-value < 0.05). Representative images of each structure are shown in the lower panel; scale bar represents 10 μm. **(b)** Representative images of appressoria CFW glucan staining and AM1::GFP in SG200 and Δ*row1* mutant strains (left panel). Scale bar represents 10 μm. The quantification of the CFW signal (lower panel) was performed using the same method as in Figure 4. The total number of appressoria is indicated above each column. Error bars represent the standard deviation from two independent replicates. The Student’s t-test statistical analysis was performed for each mutant versus the WT strain (ns, no significant; \**p*-value < 0.05).

### The **Δ***row1* mutant exhibits impaired secretion

As the *U. maydis* appressorium does not provide mechanical force for physical penetration, secretion of other proteins is essential for proper penetration: PCWDEs break down the host cell wall, while effectors manipulate host cell physiology and promote fungal penetration, colonization, and tumour formation (Lanver *et al*., 2017). Since the Δ*row1* mutant strain has poorer appressorium formation and progression inside plant tissues, and the cell wall is altered, we hypothesized that this mutant may has altered secretion, thereby compromising appressorium biology.

To investigate the potential impact of row1 deletion on secretion, we analyzed the secretomes of a WT and the Δ*row1* mutant in pathogenic filamentous growth conditions by Quantitative Mass Spectrometry. We observed an altered secretion profile for the Δ*row1* mutant, with a decrease in secretion as the main variation. We identified 39 proteins for which the difference in secretion levels between the two strains was statistically significant: 35 of the proteins were secreted less in the Δ*row1* mutant, while four were secreted more (Fig. 6a upper left panel and Table S4). Among the 35 proteins that were secreted less in the Δ*row1* mutant, we identified several membrane-related transported proteins, such as the putative vacuolar ATP synthase subunit E, an ABC transporter-domain containing protein, and an acyl-CoA binding domain (ACB) protein, which represent one of the main targets of unconventional secretion pathways (Ponpuak *et al*., 2015). Two of the 35 proteins were related to lipids: an annexin and the Scp2 effector identified in peroxisomes in *U. maydis* (Krombach *et al*., 2018). This group of proteins also included the protein Snf7 of the ESCRT III complex, which is involved in vesicle trafficking; one septin protein; a GH16 glucanase, which is involved in cell wall modification; and Row1 itself, confirming our previous suspicions about its secretion. Finally, this group of 35 proteins included several proteins associated with mitochondria and three ribosomal subunits (Fig. 6a lower left panel, Table S4). Interestingly, most of the categories in which the differentially secreted proteins were classified have been associated with unconventional protein secretion (UPS) through extracellular vesicles (EVs) (Rutter *et al*., 2022) (Fig. 6a lower left panel, Table S4).

**Fig. 6.**
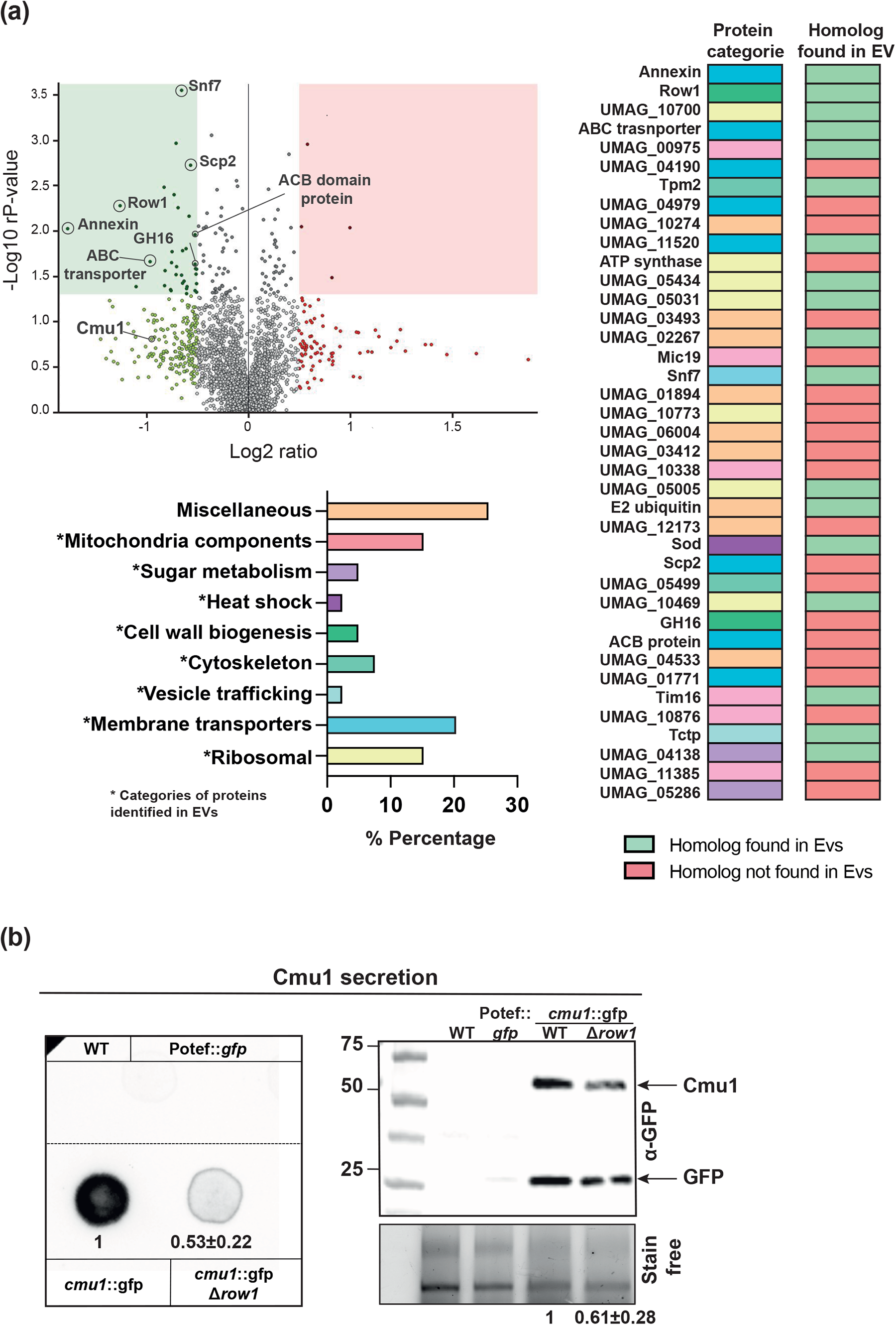
Row1 affects secretion. **(a)** Volcano plot of all proteins identified in the secretome of *U. maydis* after inducing filament conditions for 5 h. Proteins secreted less in the Δ*row1* mutant than in the WT are shown in green. The proteins represented by dark green points inside the green rectangle are differentially secreted proteins (fold change between -0.5 and 0.5, log2FC, and p-value ≤ 0.05). Proteins secreted more in the Δ*row1* mutant than in the WT are shown in red. The proteins represented by dark red points inside the red rectangle are differentially secreted proteins (fold change between – 0.5 and 0.5, log2FC, and p-value ≤ 0.05). The percentage of these proteins in each established category is shown in the lower left panel. Right panel: The presence (green) or absence (red) of homologues identified in previously characterized and purified EVs from other organisms for each of the identified proteins**. (b)** Colony secretion assay of effector Cmu1::GFP in pathogenic conditions using PD-charcoal plates. SG200 filaments expressing cytoplasmic GFP under control of the constitutive *otef* promoter served as a cell lysis control. The data show a single representative experiment out of three repeats, and quantifications are the averages and standard deviation of the mutant GFP signal relative to WT from three independent experiments. The western blot assay shows the secreted protein fraction of Cmu1::GFP in WT and Δ*row1* mutant backgrounds extracted from cells in axenic conditions. The data shown here is representative of a minimum of three repetitions, and the quantifications represent the average and standard deviation of the GFP signal relative to the stain-free control from three independent experiments.

To investigate if these proteins were found specifically in EVs, we searched for homologous proteins in pathogenic fungi in which EVs and their components have already been purified, such as *Fusarium oxysporum*, *Candida albicans*, *Cryptococcus neoformans* and *Histoplasma capsulatum*, as well as in the yeast *Saccharomyces cerevisiae* (Albuquerque *et al*., 2008; Vargas *et al*., 2015; Zhao *et al*., 2019; Garcia-Ceron *et al*., 2021). Our analysis revealed that around 50% of the 35 proteins that were secreted less in the Δ*row1* mutant had homologues in the EVs of these fungi (Fig. 6a right panel, Table S4). Although annexin did not have a homologue in these fungi, it is found in the EVs of mammals (Rutter *et al*., 2022). Curiously, we identified a homologue of Row1 named MP88 as one of the most abundant proteins in *C. neoformans* EVs (Rizzo *et al*., 2021). This suggests a function for these proteins in EV-mediated processes and indicates that Row1 might play a role in this type of secretion.

We also found in the proteomics analysis that the effector protein chorismutase 1 (Cmu1) (Djamei *et al*., 2011) was secreted less in the Δ*row1* mutant than in the WT, although the difference was not statistically significant. As Cmu1 and other effectors are not expected to be secreted by the UPS pathway, we examined its secretion by colony secretion assays under pathogenic conditions. Our results confirmed a decrease in Cmu1 secretion in the Δ*row1* mutant compared with the WT (Fig. 6b). To validate this finding, we isolated the secreted protein fraction and studied the Cmu1 amount by western blotting. In agreement with our previous observations, we found a lower amount of Cmu1 in the Δ*row1* mutant compared to the WT strain (Fig. 6b).

### Row1 is part of a fungal protein family with roles in *U. maydis* virulence

Despite the proposed role for Row1 in crucial stages of *U. maydis* pathogenic development such as appressorium formation and secretion, the loss of Row1 does not lead to a massive reduction in virulence capacity, as Δ*row1* infections still cause tumour formation (Fig. 1a). One plausible explanation is the potential presence of Row1 paralogues in *U. maydis*. Thus, we searched for proteins containing a similar sequence to Row1 using BlastP analysis. We found five putative paralogues of Row1: Umag_00961 (Row2), Umag_02921 (Row3), Umag_10474 (Row4), Umag_06162 (Row5), and Umag_03349 (Row6) (Fig. 7a, Fig. S6 and Table S5). The genes for all five paralogues are located on different chromosomes and have a similar length, and the proteins all contain a signal peptide, a serine-rich domain and O-and/or N-glycosylation sites. Row1, Row4, Row5, and Row6 have a transmembrane region, and Row5 and Row6 have a GPI-anchor site similar to that of Row1. Row2, Row3, and Row5 are predicted to be effector and apoplastic proteins (Fig. 7a, Table S6). MAFFT multiple alignment showed high conservation between all the paralogues, mostly in the central region (approximately amino acids 100–300), and the Sma3 tool associated Row2–5 with the same GO annotations as Row1 (Fig. S6, Table S6).

**Fig. 7.**
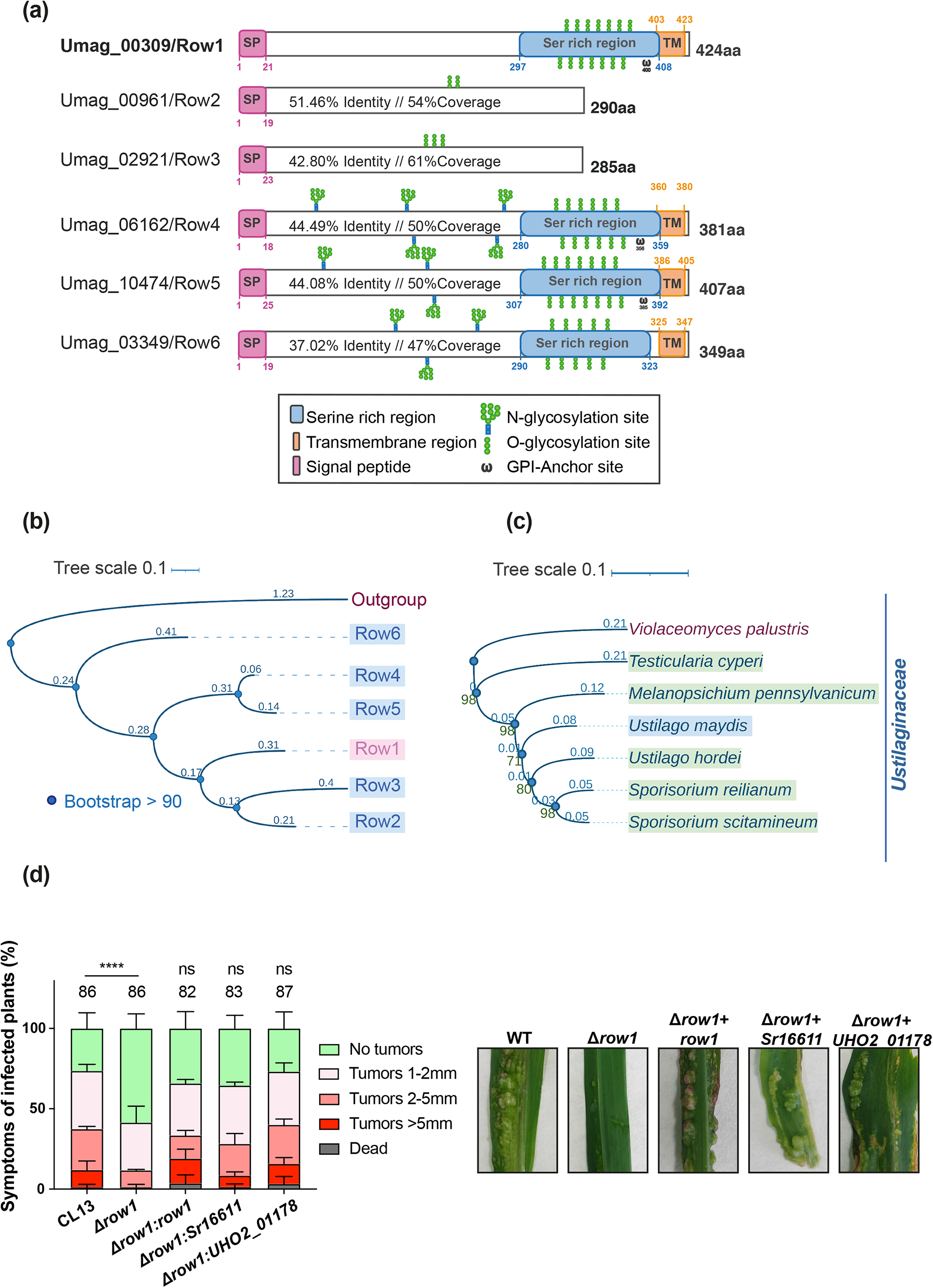
Row1 has five paralogues and is functionally conserved in the related smut pathogens. **(a)** Schematic picture of Row1 paralogues in the fungus *U. maydis*. **(b)** Phylogenetic study of Row1, Row2, Row3, Row4, Row5 and Row6. Black circles at nodes indicate bootstraps higher than 90, and distances are indicated in blue above each branch. Umag_02751 was used as an outgroup. BlastP was used to search for Row1 homologous sequences in *U. maydis*. The alignments were obtained using MAFFT v7. Phylogenetic analysis was inferred by using the maximum likelihood method. The phylogenetic trees were generated using Archaeopteryx.js and edited in iTOL. **(c)** Phylogenetic study of Row1 orthologues in the smut fungi Ustilaginalaceae family including *Sporisorium reilianum*, *S. scitamineum*, *Ustilago hordei*, *U. maydis*, *Melanopsichium pennsylvanicum*, *Testicularia cyperi* and *Violaceomyces palustris* as an outgroup. Bootstraps are indicated in green numbers down the nodes, and the nodes and distances are indicated in blue above each branch. **(d)** Left panel: Pathogenicity assay of *U. maydis* Δ*row1* mutant complemented with *S. reilianum* and *U. hordei* orthologues. Quantification of symptoms for plants infected with indicated strains at 14 dpi. The total number of infected plants is indicated above each column. Error bars represent the standard deviation from two independent replicates. The Mann–Whitney statistical test was performed for each mutant versus corresponding WT strain (ns, not significant; \*\*\**p*-value < 0.005). Right panel: Representative images of the most prevalent tumour category.

To study if members of this family have conserved functional domains, which would suggest similar roles, we conducted a structural prediction analysis using AlphaFold. The predicted structures of Row1 and all five paralogues showed a main domain corresponding to amino acids 100–300 that exhibited a high degree of superposition among all the paralogues (Fig. S7a) except for Row6, which displayed the most significant differences (Fig. S7b). To further explore the possibility that all these proteins represent a protein family, we developed a phylogenetic study. All members showed a common ancestor and small distance were represented between the five members. While Row2, Row3, Row4, and Row5 were the most related sequences and were grouped in the same clade, Row6 was placed on a separate branch (Fig. 7b). All these findings led us to conclude that these genes are part of a gene family, which we call the Row family.

To determine whether Row1 and the other members of the Row family are conserved in other smut fungi or are specific to *U. maydis*, we first searched for orthologues of Row1 in Ustilaginaceae (Zuo *et al*., 2019). A homology search and phylogenetic tree analysis, which included most representative smut fungi, revealed the presence of Row1 homologues in all available genomes of smut fungi (Fig. 7c, Table S7). To explore whether these orthologues are also functionally conserved, we complemented the *U. maydis row1* deletion mutant by introducing orthologues from *Sporisorium reilianum* and *Ustilago hordei* under the *row1* promoter of *U. maydis*. Both orthologues could fully restore the virulence phenotype of the *row1* deletion mutant (Fig. 7d).

After experimentally verifying that Row1 was conserved in Ustilaginales, we extended the study to the rest of the Row family. The phylogenetic tree grouped the six members of the family into six independent clades with a common ancestor (Fig. S8). While we observed some variability between family members, such as Row3 not being conserved in *Melanopsichium pennsilvanicum*, *Testicularia cyperi*, and *Kalmanozyma brasiliensis*, Row6 not being conserved in *T. cyperi*, and Row4 and Row5 not being present in *Moesziomyces aphidis*, the overall protein family is conserved within the Ustilaginaceae clade (Table S8 and Table S9). Next, we carried out a full conservation study of the Row protein family across fungi. We observed conservation specifically in the Basidiomycota division (Fig. 8a, Table S10). We must highlight the presence of Row family members in rust fungi, which belong to Pucciniomycotina such us *Melampsora* spp., *Phakopsora pachyrhizi* and *Puccinia* spp. (Fisher et al., 2012; Yang, 2022), members of the genus *Microbotryum* that infect flowers of different plants, and *Mixia osmundae*, a fern parasite (MIX, 1947). In addition to being found in Ustilaginales fungi (Fig. S8 and Fig. 7c), the Row family also appears in Ustilaginomycotina species belonging to the Exobasidiomycetes order, such as *Tilletia* spp., which infect wheat and triticale (Bishnoi *et al*., 2020). The Row family is also conserved in animal and human pathogenic fungi such as *Malassezia* and *Cryptococcus* spp. (Heitman, 2011), where several members of the family have been partially characterized. In *C. neoformans*, CNAG_00776 and CNAG_6000 showed similarity mainly to Row5 and Row4 (Table S11), and their deletion resulted in growth defects (Snelders *et al*., 2022) and capsule formation problems (Han *et al*., 2020), respectively. CNAG_05312, which showed similarity mostly to Row2 (Table S11), was associated with melanin granules in the *Cryptococcus* capsule (Camacho *et al*., 2019) and was identified in a PKA1 protein-induced screening, which is involved in the synthesis of the cell wall (Geddes *et al*., 2015).

**Fig. 8.**
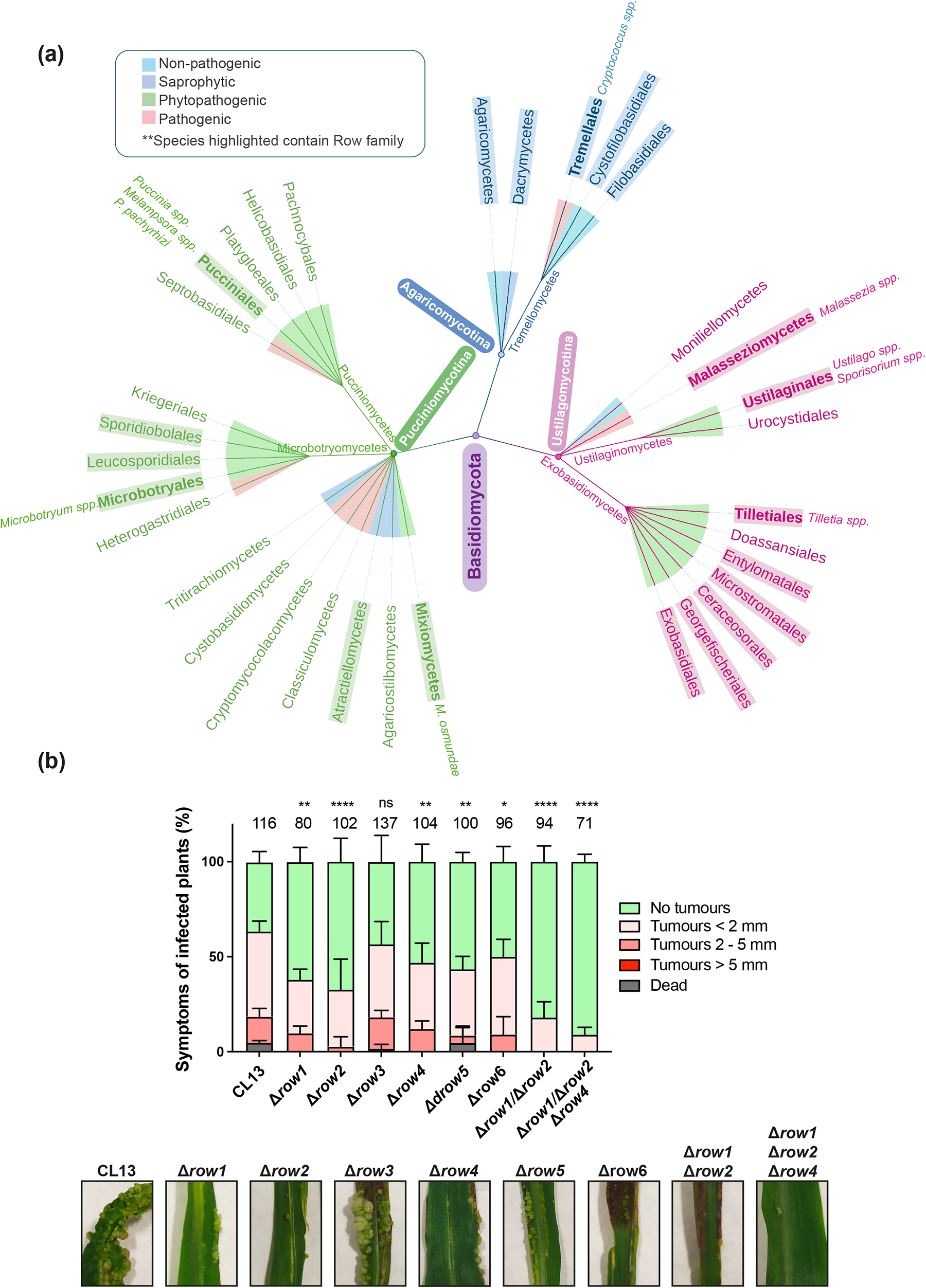
The Row family is mostly conserved in pathogenic fungi and is involved in *U. maydis* virulence. **(a)** The taxonomic tree of the Basidiomycota clade illustrates the conservation of the Row family across fungi. The Basidiomycota clade (purple) comprises three subdivisions: Agaricomycotina (blue), Pucciniomycotina (green), and Ustilagomycotina (pink), represented by nodes. Fungi belonging to these subdivisions are classified into four groups: non-pathogenic (cyan), saprophytic (blue), phytopathogenic (green), and animal pathogenic (pink). The highlighted genera include members of the Row family, and examples of species in which the family is conserved are written next to them. The taxonomic tree was obtained using the National Center for Biotechnology Information taxonomy browser. **(b)** Quantification of symptoms for plants infected with mutants Δ*row1*, Δ*row2,* Δ*row3,* Δ*row4,* Δ*row5,* Δ*row6*, the double mutant Δ*row1*Δrow2 or the triple mutant Δ*row1*Δ*row2*Δ*row4* at 14 dpi. The total number of infected plants is indicated above each column. Error bars represent the standard deviation from three independent replicates. The Mann–Whitney statistical test was performed for each mutant versus the WT strain (ns, not significant \**p*-value < 0.05, \*\**p*-value < 0.005, \*\*\*\**p*-value < 0.0001). Representative images of the most prevalent tumour category for WT and mutant strains are shown in the lower panel.

The observed conservation of this protein family in pathogenic fungi, coupled with our findings, suggests that Row1 is part of a family of proteins that may have similar virulence-related functions. To scrutinize this hypothesis, we studied the potential role of the Row family in infection by deleting each gene in the CL13 background of *U. maydis.* While we found no significant differences in stress responses (Fig. S3), all mutants except Δ*row3* had reduced virulence compared to the WT, exhibiting a phenotype similar to Δ*row1* (Fig. 8b and Fig. S9). Of all mutants, Δ*row2* presented the lowest virulence capacity. As Δ*row2* and Δ*row4* had the lowest capacity for infection, and the expression profile of these genes during the first stages of infection is similar to that of *row1* (Fig. S10), we considered the possibility that these proteins have similar functions during pathogenic development. We found that the Δ*row1*Δ*row2* double mutant caused significantly less severe symptoms in infected maize plants than did single mutants. The triple mutant Δ*row1*Δ*row2*Δ*row4* even showed less severe symptoms than the double mutant (Fig. 8b), indicating that Row1, Row2 and Row4 might have redundant functions during pathogenesis. These findings support the hypothesis that Row1 and its homologues may have similar functions in pathogenic development.

### Row2 has a similar localization to Row1 and is also important for secretion and cell wall modification

To further test the idea of similar functions across the Row family, we selected Row2, which showed the highest virulence defect, to analyse cellular localization and its possible role in secretion and cell wall structure. We observed a similar location for Row2 and Row1, with predominant signals at the ER and plasma membrane and an additional slight accumulation in the nucleus (Fig. 9a). Based on the predicted signal peptide and its localization, we hypothesized that Row2 could be part of the secretory pathway. We carried out a colony secretion assay that confirmed Row2 secretion (Fig. 9b). We then investigated if Row2 also plays a role in effector secretion. We observed a decrease of the Cmu1 secreted fraction in the Δ*row2* mutant compared to WT, suggesting that Row2 is required for efficient Cmu1 secretion (Fig. 9c). Since Row1 and Row2 share similar localization and both affect secretion, we investigated whether they have redundant functions in the pathogenic process. We introduced an extra copy of *row2* in the Δ*row1* mutant, thereby compensating for the loss of *row1*, which rescued the pathogenic defects of the mutant (Fig. 9d). This suggests a functional redundancy between the two proteins, indicating that they may have overlapping roles in pathogenesis.

**Fig. 9.**
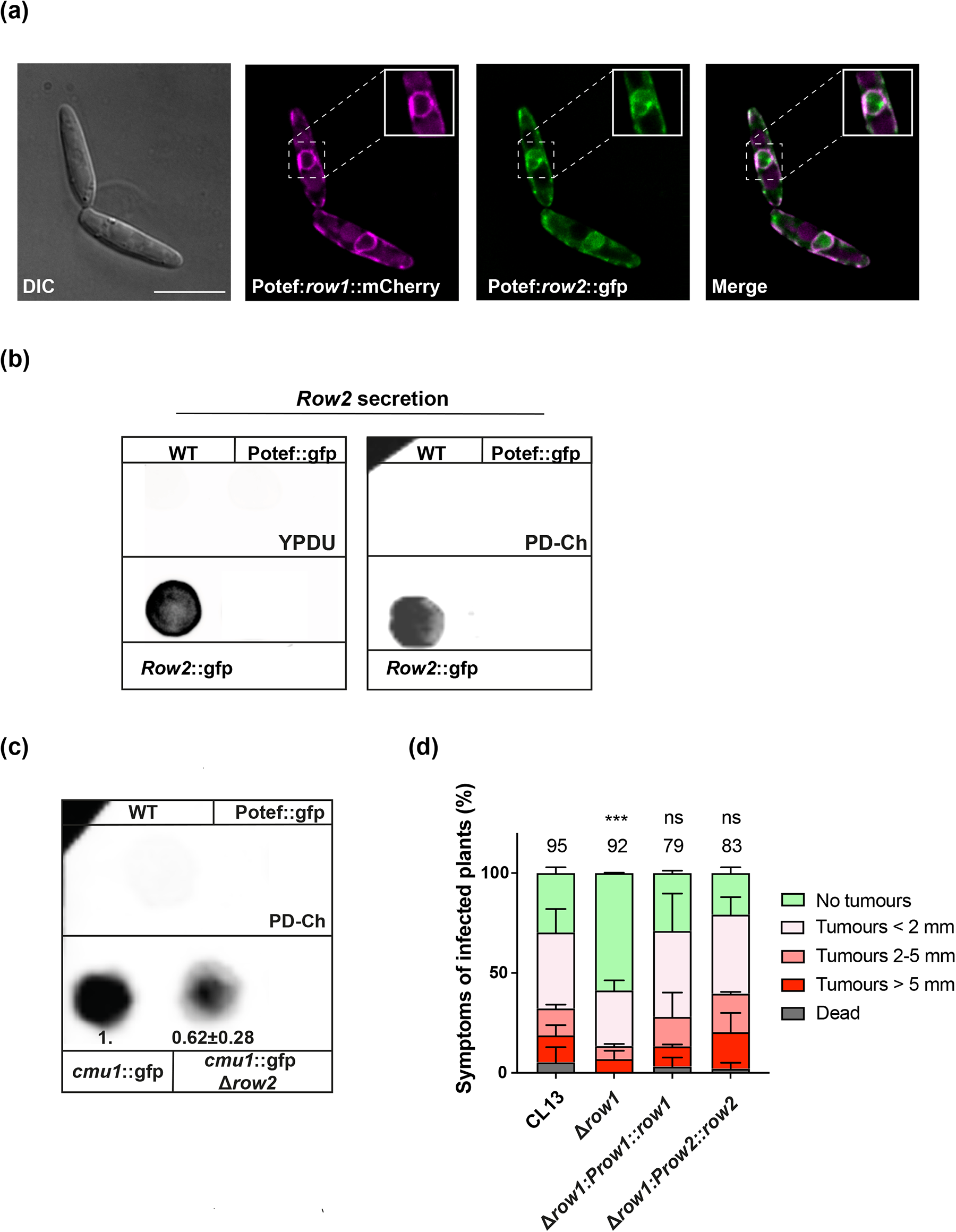
Row2 is a secreted protein with a similar localization to Row1 and is involved in secretion. **(a)** Row1::mCherry co-localization with Row2::GFP in growing cells. Scale bar represents 10 μm. **(b)** Colony secretion assay of Row2::GFP in non-pathogenic and pathogenic conditions using YPDU and PD-charcoal plates. The SG200 WT strain and SG200 cells expressing cytoplasmic GFP under the control of the constitutive *otef* promoter served as cell lysis controls. **(c)** Colony secretion assay of Cmu1::GFP in WT and Δ*row2* mutant backgrounds performed under pathogenic conditions using PD-charcoal plates. The data show a single representative experiment out of at least three repeats, and quantifications are the averages and standard deviation of the mutant GFP signal relative to WT from at least three independent experiments. **(d)** Infection assay of *U. maydis* Δ*row1* mutant complemented with Row2 homologue at 14 dpi. The total number of infected plants is indicated above each column. Two biological replicates were analysed. The Mann–Whitney statistical test was performed for each mutant versus the WT strain (ns, not significant; \*\*\**p*-value < 0.0005).

Based on these findings and considering that the double mutant had less capability for infection than the single mutant (Fig. 9b), we generated a Δ*row1*Δ*row2* double mutant in the AB33 strain to explore whether it would also display defects in cell wall composition or structure. Analysis with CFW suggest an effect of Row2 on glucans composition (Fig. 3b), and electron microscopy showed that this mutant had a thicker inner layer than WT and Δ*row1* single mutant (Fig. 4a). We also observed a decrease in WGA chitin intensity in the double mutant when compared with both the WT and Δ*row1* mutant strains (Fig. 3a), which suggests that a thicker cell wall could impede access of the stain to the chitin. These findings strongly indicate the involvement of Row1 and Row2 in fungal cell wall remodelling and suggest a cooperative relationship between the two proteins in this process.

## DISCUSSION

Our findings highlight Row1 as important for appressorium cell wall remodelling and protein secretion. We show that Row1 belongs to a conserved family of proteins that are involved in virulence, are found predominantly in pathogenic fungi of the Basidiomycota clade and have potential functional similarities with Row1.

### The role of Row1 in the infection process

As Row1 is expressed mainly on appressoria, and its absence leads to defects in appressoria formation and penetration and to alterations in the cell wall, it is tempting to think that the defect in the cell wall observed in the Δ*row*1 mutant may cause appressoria penetration problems. In contrast to fungi with melanized appressoria such as *M. oryzae*, fungi with non-melanized appressoria, such as *U. maydis*, rely on PCWDEs and effector secretion to break the plant cuticle and establish pathogenic development (Kubicek *et al*., 2014; Lanver *et al*., 2014; Chethana *et al*., 2021; Bradley *et al*., 2022). Thus, the thicker cell wall observed in the Δ*row1* mutant may cause some defect in secretion that leads to defective penetration. In agreement with this idea, we found that the Δ*row1* mutant had a defect in the secretion of some important proteins for infection than WT, including effectors such as Cmu1 and Scp2 (Table S4 and Fig. 6). This correlation between cell wall thickness and secretion has been already observed in the pathogenic fungi *Aspergillus nidulans*, where a weak cell wall has been suggested as the possible cause of higher secretion (Boppidi *et al*., 2018). In addition, a defective or absent cell wall has been reported as affecting secretion in *Neurospora crassa* and *Aspergillus nidulans*, respectively, which supports the role of the fungal cell wall in secretion (Peberdy, 1994; Sietsma *et al*., 1997). In this scenario, Row1 may have an active role in fungal cell wall modification, potentially acting as a catalytic enzyme. Although most remodelling enzymes have annotated domains that involve carbohydrate binding or enzymatic activity, neither Row1 nor the rest of the Row family members have any annotated domain (Fig. 2a). However, this has also been observed in proteins such as the effector Sta1 in *U. maydis* (Tanaka *et al*., 2020) and the GPI proteins Pga13 and Pga31 in *C. albicans* (Plaine *et al*., 2008; Gelis *et al*., 2012), which lack annotated glycoside hydrolases or carbohydrate-binding domains but have been suggested to have an active role in cell wall processes. Furthermore, as we have shown here, Row1 has sequence and structural homology to some proteins of species in Agaricomycotina (Fig. 8a). Some of these proteins, mostly in species in the Agaricales order, were annotated as carbohydrate-binding module family proteins due to the presence of the CBM13 domain, which is commonly found in enzymes involved in the degradation of complex carbohydrates (Fujimoto, 2013) such as glycosyl hydrolases (Boraston *et al*., 2000; Notenboom *et al*., 2002)). However, the similarity between Row1 and these proteins did not include the region corresponding to the CBM13 domain (Fig. S11), which could have been lost in other Basidiomycota organisms or acquired later by Agaricales fungi. We speculate that like these proteins, Row1 could play a role in cell wall modification, potentially contributing to enzymatic activity independently of the CBM13 domain. Supporting this hypothesis, Sma3 analysis predicted a putative involvement of Row1 in polysaccharide catabolic processes and hydrolase activity on glycosyl bonds. As Δ*row1* mutant filaments had a thicker inner cell wall (Fig. 4), higher resistance to glucan degradation (Fig. 3c), increased CFW accumulation in the tip of the filament and during appressoria formation (Fig. 3b and 5b), but no chitin defects in WGA staining (Fig. 3a), we speculate that Row1 is involved in glucan modification during appressoria formation.

Nevertheless, as secretory pathways contribute to the structure of cell walls and host interactions (Latgé, 2007), another possibility that cannot be ruled out is that Row1 has a role in the secretion process, thereby altering fungal cell wall architecture. While the classical ER/Golgi-dependent pathway is responsible for the secretion of most extracellular proteins, many proteins without a signal peptide follow either a vesicle-independent or vesicle-dependent UPS route (Rabouille, 2017; Dimou & Nickel, 2018). In the latter case, proteins can be released within EVs, which are small lipid-bilayer compartments involved in transporting proteins, lipids, nucleic acids, and other macromolecules outside the cell (Shoji *et al*., 2014). These EVs are formed inside late endosomes as intraluminal vesicles and then released when the late endosome fuses with the plasma membrane (van Niel *et al*., 2018; Vats & Galli, 2022). We show that Row1 exhibits vesicular movement, accumulating mainly at the tip of the hyphae, with partial co-localization with Yup1, a t-SNARE present in endosomes (Fig. 2d). These observations and the specific pool of proteins that are secreted less in the Δ*row1* mutant than in WT, rather than a general secretion problem (Fig. 6a), lead us to propose a potential role for Row1 in these vesicular processes as an alternative possibility. In agreement with this proposal, our mass spectrometry analysis showed less secretion of EV-related proteins than the WT strain (Fig. 6a, Table S4). In addition, a significant percentage of the differentially secreted proteins are mitochondrial components. It has been demonstrated in mammals that cells selectively regulate the packaging of mitochondrial protein into EVs to prevent the release of damaged components that would otherwise act as pro-inflammatory damage-associated molecular patterns (Todkar *et al*., 2021). Mesenchymal stem cells also undergo mitophagy in response to oxidative stress, packaging mitochondrial components in EVs for cellular transfer (Phinney *et al*., 2015). All these data suggest that Row1 may be involved in UPS through EVs. The alteration in the secretion of these EVs would easily explain other effects of the Δ*row1* mutant, such as cell wall and appressoria penetration defects. EVs have been proposed to be involved in the remodelling of the cell wall to facilitate their transit across it by carrying wall-remodelling enzymes as part of their cargo, such as β-glucosidases, chitin-deacetylases or endochitinases (Rodrigues *et al*., 2007; Albuquerque *et al*., 2008; Oliveira *et al*., 2010; Brown *et al*., 2015). In our analysis, we detected lower levels of the GH16 glucanase and chitin-deacetylase 5 in the Δ*row1* mutant than in the WT, although the difference in the latter was not statistically significant. We also found a decrease in the Δ*row1* secretion of the effector Scp2, which is essential for proper appressorium formation (Krombach *et al*., 2018), and annexin, which is associated with the cell wall in the fungus *Phytophthora infestans* and plays a crucial role in the penetration of this pathogen into the host tissue (Grenville-Briggs et al., 2010). In addition, we detected a defect in the Δ*row1* secretion of the effector Cmu1 (Fig. 6b), which should be secreted through the conventional secreted pathway. A recent study demonstrates that effectors expressed during the infection of corn are contained within the EVs of *Fusarium graminearum* (Garcia-Ceron *et al*., 2021). This finding raises the possibility that Cmu1 could also be present in unconventional secretion pathways or be indirectly affected by the role of Row1 in UPS. In *C. neoformans*, one of the major components of EV membranes is the protein MP88, which is homologous with Row1 (Rizzo *et al*., 2021) (Table S11), which suggest a direct role for Row1 in the proper formation or maturation of EVs.

We have proposed two alternative scenarios regarding the potential function of Row1: as a cell wall remodelling enzyme affecting secretion, or as a protein involved in secretory pathways that affect the cell wall. However, these scenarios are not mutually exclusive. Thus, we propose a third scenario that encompasses both hypotheses, in which Row1 may be a component of EVs that facilitates glucan degradation for the cell wall remodelling that is necessary for the proper secretion of EVs. In the absence of Row1, EVs are unable to efficiently remodel the cell wall, leading to defects in their own secretion. Since the components of EVs are crucial for pathogenesis (GarciaCeron:2021bt, Albuquerque *et al*., 2008; Vargas *et al*., 2015), a reduction in their secretion could result in deficiencies in pathogenesis, particularly during the penetration stage.

### Functions of the Row family

We have shown that Row1 belongs to a family of six Row proteins conserved in Basidiomycota. All of these proteins except for Row3 have roles in infection in *U. maydis* (Fig. 8b and Fig. S9). The conservation of the globular central domain, which contains the possible glucan catabolic activity, is consistent with the idea that these proteins all share a main role in cell wall remodelling. However, the similar membrane localization and secretion defects observed for Row1 and Row2 indicate that a secretion role for family members cannot be discounted. Our findings that phylogenetic analysis demonstrates a common ancestor and that Row2 compensates for Δ*row1* defects in tumour formation reinforce the idea of a shared main function for Row family members (Fig. S8 and Fig. 9c). However, although they may share a main role, each protein would have evolved to play a different specialized function during infection, likely at different moments of infection. This is supported by the diverse expression pattern during pathogenic development (Fig. S10): Row1 and Row2 are expressed at the first stages, followed by Row3 and Row4, then Row6 during biotrophic establishment, and finally Row5 when tumorigenesis begins (Fig. S10). Previous studies have exemplified protein functional specialization in different fungal systems. For instance, a family of three ferroxidases involved in iron uptake in *Mucor circinelloides* are differentially expressed in yeast and hyphae forms (Navarro-Mendoza *et al*., 2018). Cerato-platanins, small cysteine-rich fungal secreted proteins (Pazzagli *et al*., 1999), have crucial roles in various stages of the host-fungus interaction process and present distinct expression profiles during the life cycle of different pathogen fungi (de O Barsottini *et al*., 2013, Gaderer et al., 2014), similar to that observed for Row members.

Overall, we present here a new family of conserved proteins, the Row family, with important roles in infection that may share a common function, probably in cell wall remodelling or as UPS proteins. The Row family may represent a new group of target proteins for the development of antifungal compound with a wide spectrum due to the high conservation they have on pathogenic fungi.

## Acknowledgements

We would like to thank the Genetics Department for their useful discussions and comments. Victor Manuel Carranco, Sandra Romero, Blanca Navarrete and Adrián Prieto for the technical assistant. Cristina Vaquero Aguilar from CITIUS (Universidad de Sevilla) for technical support with Electron Microscopy. This research was supported by MCIN/AEI/10.13039/501100011033/ and by “ERDF A way of making Europe”, grant number BIO2016-80180-P and MCIN/AEI/10.13039/501100011033/ grant number PID2019-110477GB-I00 to JII.

## Data availability

The data that support the findings of this study are openly available in (repository name and URL will be available after acceptance), reference number (reference number will be available after acceptance), and in the supplementary material of this article.

## Competing interest

The authors declare no competing interests.

## Author contributions

MD.P-O, J.I.I and R.R.B planned and designed the research. MD.P-O generate strains, performed the experiments, and analyzed the data. L.T.G performed Mass Spectrometry experimental procedures. MD.P-O and R.R.B wrote the original manuscript with input from all coauthors.

## Notes

### Competing Interest Statement

The authors have declared no competing interest.

