## Supplemental Material for "Row1, a member of a new family of conserved fungal proteins involved in infection, is required for appressoria functionality in Ustilago maydis"

### New Phytologist Supporting Information

**The following Supporting Information is available for this article:**

**Methods S1** Plasmid cloning and strain generation; Strain growth conditions and virulence assay in maize; Microscopy analysis, sample preparation, and microscope characteristics and settings; Sample preparation for proteomic assay; Sequence alignment, phylogenetic analysis and predictive analysis tool.

**Fig. S1** Δ*row1* affects virulence in the genetic backgrounds FB1xFB2, CL13 and SG200.

**Fig. S2** Lack of Row1 does not affect growth, cell length or adhesion.

**Fig. S3** Δ*row*1–6 mutants do not present cell growth defects under saline, oxidative, cell wall or reticular stresses.

**Fig. S4** Row1 could be a secreted protein.

**Fig. S5** Δ*row1* mutant does not exhibit defects in filament length or morphology.

**Fig. S6** Row family sequence alignment.

**Fig. S7** Row family structural alignment.

**Fig. S8.** Row family is conserved in Ustilaginales.

**Fig S9** Row family members are involved in *U. maydis* virulence.

**Fig. S10** Row family members are differentially expressed during infection.

**Fig. S11** Row1 does not have a CBM13 domain, which is conserved in homologues belonging to the Agaricales order.

**Table S1** Strains used in this study.

**Table S2** Plasmids used in this study.

**Table S3** Accession numbers.

**Table S4** Proteins that are differentially secreted in the Δ*row1* mutant and WT strains, and their homologues identified in EVs (separate Excel file).

**Table S5** Protein homologues of Row1 in *U. maydis.*

**Table S6** Characteristics of Row family proteins (separate Excel file)

**Table S7** Protein homologues of Row1 in Ustilaginaceae.

**Table S8** Row family conservation in Ustilaginaceae.

**Table S9** Protein homologues of Row family members in Ustilaginaceae.

**Table S10** Protein homologues of Row family members in Basidiomycota (separate Excel file).

**Table S11** Homologues of Row1 in *Cryptococcus neoformans.*

#### Methods S1:

##### Plasmid cloning and strain generation

*Escherichia coli* DH5α and *pJET1.2/blunt* (ThermoFisher Scientific) were used for cloning purposes. Growth conditions for *E. coli* (J *et al.*, 1989) and *U. maydis* (Gillissen *et al.*, 1992; Brachmann *et al.*, 2004) have been previously described.

NEBuilder® HiFi DNA Assembly (BioLabs) or standard molecular cloning techniques were used for plasmid construction (J *et al.*, 1989). An adjusted version of *U. maydis* DNA isolation and transformation procedures were carried out following the protocol of (Bösch *et al.*, 2016).

For the deletion of *row1, row2 and row6* deletion strains, 1 kb fragments of the 5' and 3' flanks of the ORF from the gene of interest (*goi*) were generated by PCR using Q5 High-Fidelity DNA polymerase (New England Biolabs) amplified from CL13 genomic DNA, using the primers goiKO5-1 and goiKO5-2 (containing a SfiI restriction site) to amplify the 5' flank and goiKO3-1 (containing a SfiI restriction site) and goiKO3-2 to amplify the 3' flank. The PCR products were digested with SfiI and ligated with the 1.9-kb SfiI carboxin, 1.4-kb SfiI noursethricin (ClonNAT), 2-kb SfiI geneticin or 1.9-kb SfiI hygromicin resistance cassettes (Brachmann *et al.*, 2004). Constructs were cloned into pJET1.2/blunt (ThermoFisher Scientific) plasmids to generate the plasmids pJET Δ*row1*:NatR, pJET Δ*row2*:hHygR and pJET Δ*row6*:CbxR . Each plasmid was amplified by PCR using the primers goiKO5-1/goiKO3-2 to transform different *U. maydis* strains by homolog recombination.

Deletion of *row3, row4 and row5* was generated by assembling of the 1 kb PCR fragments corresponding to the 5' and 3' flanking regions of the ORF of interest, using the primers goiKO5-1 and goiKO5-2, the resistance cassettes,using the Resistance_Fwd_goi and Resistance_Rev_goi primers, and the PBSK plasmid, previously digested with BamHI, using the NEBuilder® HiFi DNA Assembly Master Mix (BioLabs) to generate the plasmids pBSK Δ*row3*:GenR, pBSK Δ*row4*:CbxR, and pBSK Δ*row5*:HygR. The deletion cassettes were obtained by PCR of the generated plasmids using the primers goiKO5-1/goiKO3-2 and transformed in the *U. maydis* strains. The triple deletion strain CL13 Δ*row2*/Δ*row1*/Δ*row4* was generated as follows. The *row1* deletion cassette was amplified from (pJET Δ*row1*:NatR) and transformed into the strain CL13Δ*row2*. After verification of the double deletion strain, the Δ*row4* deletion construct was amplified from pBSK Δ*row4*:CbxR plasmid using the primers Ko5-1 *Row4*_PCR / Ko3-2 Row4_PCR and transformed into CL13*/*Δ*Row2/* Δ*row1*. The double deletion strain AB33 Δ*row1/*Δ*row2* was generated by the amplification of Δ*row2*:CbxR using the primers Ko5-1 *Row2*_PCR / Ko3-2 Row2_PCR and transformed into AB33 Δ*row1*.

To complement the strain CL13Δ*row1*, the construct p123 P*row1*:*row1* was generated. Genomic DNA from CL13 containing promoter and ORF of *row1* was amplified by PCR with primers Fwd-Row1-Promoter (PuvII) and Rev-Row1 (NotI) and digested with PvuII and NotI. The fragment was ligated into the plasmid p123 previously digested with both restriction enzymes using T4 DNA ligase (New England Biolabs). To generate AB33 P*row1*:row1::gfp the construct p123 P*row1*:*row1*::gfp was generated. Procedures were the same as p123 P*row1*:*row1* plasmid construction but the promoter and open reading frame of *row1* fragment were amplified using primers Fwd-Row1 Promoter (PuvII) / Rev-Row1 (NcoI) and digested with PvuII and NcoI restriction enzymes and cloned into p123 plasmid digested with these enzymes. Plasmids were linearized with SspI and integrated into the AB33 ip locus by homologous recombination. To generate AB33 P*row1*:*row1*::*gfp* mrfp:HDEL, Potef-cals-mrfp-HDEL::hygR plasmid was linearized with SspI and integrated into the AB33 P*row1*:*row1*:*gfp* ip locus by homologous recombination.

To generate complementation strains of CL13Δ*row1* by *row1* orthologues from other smut pathogens, the promoter region of *U. maydis* row1 was amplified by PCR using primers Fwd-Row1_Promoter (PuvII) / Row_NcoI and cloned into a digested PvuII and NcoI p123 plasmid using T4 DNA ligase, obtaining the p123 $P_{row1}$ plasmid. Then, the open reading frame of *row1* orthologues from *S. reilianum* and *U. hordei* were amplified by PCR using primers Fwd_Sr11661_NEB / Rev_Sr116611_NEB and Fwd_UHO2_01178_NEB / Rev_UHO2_01178_NEB using genomic DNA from *S. reilianum SRZ2*, and *U. hordei* strain *4875-4* respectively. Amplified fragments were assembled using the NEBuilder® HiFi DNA Assembly kit (BioLabs) into NcoI–NotI digested p123 P*row1*::GFP. Generated plasmids were linearized with SspI and and integrated in the ip locus of CL13Δ*row1* (Loubradou *et al.*, 2001).

To introduce *row2* into the strains CL13Δ*row2* and CL13Δ*row1*, the construct p123 P*row2*:*row2* was generated. Promoter and ORF of row2 from genomic DNA from CL13 was amplified by PCR with primers Fwd-Prow2_KpnI / Rev-Prow2_NotI and digested with KpnI and NotI. The fragment was ligated into the plasmid p123 previously digested with both restriction enzymes using T4 DNA ligase (New England Biolabs). Plasmids were linearized with SspI and integrated into the CL13Δ*row2* and CL13Δ*row1* ip locus by homologous recombination.

To tag *row2* under otef promoter, p123 plasmid was digested with XmaI and NcoI restriction enzymes. *Row2* was amplified by PCR using primers Row2_XmaI_NEB / Row2_NcoI_ and cloned into digested p123 using NEBuilder® HiFi DNA Assembly Master Mix.

To obtain p123 Potef:*yup1*::mCherry and *Potef:row1*::mCherry plasmid, *yup1*, *row1* and *mCherry* were amplified using the primers Yup1-Fwd (NEB) / Yup1-Rev (NEB), Umag_00309-Fwd (NEB) / Umag_00309-Rev (NEB) by PCR from SG200 genomic DNA, and mCherry-GenR_fwd (yup1) / mCherry-GenR_rev (yup1) and mCherry-GenR_NEB_Fwd (309) / mCherry-GenR_NEB_Rev (309) from pMF5-15g plasmid. Yup1 and mCherry (yup1) and row1 and mCherry (309) were cloned into p123 plasmid digested with PvuII and NotI using NEBuilder® HiFi DNA Assembly kit (BioLabs). To generate SG200 Potef:*cmu1*::gfp, Potef:cmu1::gfp plasmid was linearized with SspI and integrated into the SG200 ip locus by homologous recombination.

**Growth conditions**

Filamentous growth of AB33 derivatives was induced by shifting cells of an exponential growing culture ($OD_{600}$ of 0.4–0.5) from YEPSL medium to nitrate minimal medium ($NO_3$-MM) supplemented with 1% glucose. Cells were incubated at 28°C shaking with 180rpm.

For charcoal mating and filamentation, cells were grown on YEPSL until exponential phase ($OD_{600}$ of 0.6-0.8), harvested via centrifugation (3000rpm, 4 min, room temperature) and washed twice with sterile distilled water. Solopathogenic strains (SG200) was diluted in $H_2O$ to a final $OD_{600}$ of 0.8 and 5 µL were potted onto Potato Dextrose (PD) – charcoal plates and grown for 24-48 hours at 25°C. Strains of compatible mating type (FB1xFB2) were mixed in a 1:1 ratio. Mating or filamentation is observed by the formation of white, fuzzy filaments. Three biological replicates with three technical replicates were performed.

For studies of growth rates and cell length and morphology, cells were grown on YEPSL to exponential phase and then diluted in the same media to an $OD_{600}$ of 0.05 and grown until exponential phase. For morphology examination cells were observed under the Inverted Microscope Olympus IX71. For growth curve assay, cells were grown until an $O_{595}$ of 1-1.2 with continuous shaking in a 96 well plates, $OD_{595}$ was monitored every 15 min with a Spark 10M (Tecan) fluorescence microplate reader coupled to a Tecan Freedom Evo 100 liquid handling platform. Three biological replicates with three technical replicates were performed.

**Virulence assay in maize**

For pathogenicity experiments, *U. maydis* strains were grown in YEPSL medium to exponential phase to an optical density $OD_{600}$ of 0.6-0.8, concentrated to $OD_{600}$ of 3 for CL13 infections, $OD_{600}$ of 1 for SG200 infections and $OD_{600}$ of 0.5 for FB1xFB2 infections. Cultures were washed twice with sterile distilled water, and injected into 7-day old maize (*Zea mays*) seedlings (Early Golden Bantam). Tumor formation was quantified 14 days post infection. At least three independent infection experiments were carried out and statistical significance was assessed using Mann-Whitney test considering significant if p values were <0.05.

**Microscopy analysis and samples preparation**

Cells for microscopy analysis were placed into an agarose pad (1,8% agarose in PBS 1X) on a microscopy slide for immobilization.

To record Row1::GFP mobility along the filaments toward the tip, spinning-disk confocal microscope. In order to observe endosome movement and Row1 colocalization, AB33 strains were cultivated in 5mL of YEPSL and grown to an $OD_{595}$ of 0.4. Filamentous growth was induced by shifting cultures to NM (1% glucose) for 4-5 h at 28℃. GFP and mCherry fluorescence were simultaneously detected using Spinning Disk Confocal Yokogawa CSUW1.

For transmission electron microscopy, 1 mL of filament cultures induced for 5 hours were centrifuged at 3000 rpm for 5 minutes. The resulting pellet was resuspended in 1 mL of 2.5% glutaraldehyde solution and incubated at 4ºC for 1 hour to fix the samples. After fixation, the samples were washed three times with 0.1 M sodium cacodylate buffer to remove excess of fixative.

*U. maydis* AMI:Cherry co-localization with Row1::GFP during penetration and hyphae proliferation inside maize leaf tissue was visualized by confocal microscopy (Leica SP5 MP-AOBS).

**Microscope characteristic and settings**

**Microscopy System DeltaVision:** Olympus IX71 inverted microscope with a range of objectives allowing for versatile magnification options. It is equipped with a CoolSnap HQ camera for high-quality imaging. The system offers environmental control capabilities, enabling precise temperature and CO2 regulation for live-cell imaging. It supports brightfield illumination and differential interference contrast (DIC). Fluorescence was detected using the follow filters: GFP491 (Excitation:470/40, Emission:528/38), DsRed561 (excitation:555/28, emission:617/73) and DAPI488 (excitation:360/40 / emission:457/50). Softworx software is used for image processing (deconvolve images).

**Spinning Disk Roper Scientific microscope system:** Olympus IX-81 inverted microscope and a Yokogawa CSU-X1 spinning disk confocal unit. For image capture, the microscope is equipped with CoolSnap HQ2 and Evolve cameras, ensuring high-quality image acquisition. Additionally, it incorporates temperature control functionality for live-cell experiments and the Bioptech FCS2 Perfusion System. The Metamorph software is utilized for data acquisition, image processing, and analysis. GFP491 (emission: 480/40) filter and Nomarsky contrast techniques for sample imaging was used.

**Spinning Disk Confocal Yokogawa CSUW1:** head with excitation lasers and filters from 3i (Intelligent Imaging Innovations). Each movie is 20 seconds (FilterQ 488q/560q, timelapse interval 250ms, laser 50 and exposition 25ms). GFP and mCherry fluorescence filters were simultaneously detected using a two-channel imager (DV2, Photometrics, Tucson, AZ). The resulting movies were converted to kymographs using ImageJ.

**Leica SP5 MP- AOBS:** inverted microscope model DMI 6000 equipped with a range of objectives. It features two PMT and two Hybrid GaAsP detectors with spectral detection, as well as galvo and resonant scanners for precise imaging. The microscope offers laser lines at 405 nm, 561 nm, 594 nm, and 633 nm, along with a multiphoton laser Mai Tai Deep See with a wavelength range from 690 nm to 1040 nm. It is equipped with two external non-descanned detectors (NDDs) and a chamber temperature controller.

**Zeiss Model Libra** 120 transmission electron microscope operate at (80 kV voltage). Images were taken in the General Research Services of the University of Seville (CITIUS).

##### Protein and blotting assays

For protein secretion assay *U. maydis* cultures were grown overnight in YEPSL medium to an $OD_{600}$ of 0.6-0.8 and harvested by centrifugation. Cultures were washed once and resuspended in 250 ml MM-$NH_4$ media containing 2% glucose and grown at 28ºC to an $OD_{600}$ of 0.6-0.8. Cells were harvested by centrifugation and the supernatant was collected and incubated at 4ºC for 30 minutes with 0.02% deoxycholate. Subsequently, proteins were precipitated following next steps: 10% trichloroacetic acid (TCA) was added, solutions were mixed by inversion, stored overnight at 4ºC, centrifuged at 12,000 rpm for 30 minutes at 4ºC and washed four times with 100% acetone. Finally, they were resuspended in TS buffer (Urea 7M, thiourea 2, CHAPS 4%) using a thermomixer (1200rpm, 10min). Protein concentration was measured using the RC DC Protein Assay kit (Bio-Rad).

For cytosolic protein extraction filaments induced for 5h and non-induced cultures were ground into a powder using a mortar/pestle under liquid nitrogen. Grounded samples were resuspended in lysis buffer (20 mM Tris-HCl, 0.5 M NaCL, pH 7.4) with protease inhibitor cocktail (cOmplete Tablets, EDTA-free, Roche) and centrifuged at 14000 rpm for 30 min at 4ºC and supernatant was collected.

60 µg of each protein fraction was separated by SDS-PAGE and detected by western blot analysis as explained (Marín-Menguiano *et al.*, 2019) Image gel and membrane acquisition was carried out with ChemiDoc MP Imaging System (Bio-Rad). Band quantification analysis was carried out by ImageLab software using stain free for protein quantity normalization. Three biological replicates were developed for each experiment.

Secretion assay was performed as previously described (Moreno-Sánchez *et al.*, 2021) using YPDU and PD-Ch media plates. Signal quantification was obtained using ImageLab program. At least, three biological replicates were developed for each experiment.

**Sample preparation for Mass Spectrometry assay**

The supernatant of 250mL of filaments induced for 5 hours were collected and precipitated with TCA as previously described. The resulting protein pellet was dried using a vacuum pump and solubilized by adding 500 µl of solubilization buffer (Urea 7M, Thiourea 2M, CHAPS 4%). Protein content was quantified using the RC-DC Protein™ Assay kit (Bio-Rad). 50 µg of secreted proteins from each culture condition were used for labelling. First, samples were adjusted to a volume of 83 µl with TrisHCl 100 mM pH7.5, reduced with TCEP 10 mM for 1 h at 55ºC and alkylated with Iodoacetamide 18 mM for 30 min in darkness. After that, proteins were precipitated o/n at -20ºC with 6 vol. acetone.

Reaction was stopped using formic acid to 0.5% and samples were labelled with the isobaric tags following manufacturer instructions, using channels 126, 127N, 127C, 128N, 128C, 129N, 129C, 130N and 130C. 5ug of every tagged sample were mixed in a single sample tube. OMIX C18 tips (Agilent Technologies) were used for concentrating and desalting tagged peptide extracts. Sample was dried and resuspended in 0.1% trifluoroacetic acid and injected in nano-HPLC system. Protein digested samples were separated in a Thermo Scientific TM Easy nLC system using a 50cm C18 Thermo Scientific™ EASY-Spray™ column. The following solvents were employed as mobile phases: Water 0.1% Formic Acid (phase A) and Acetonitrile, 20% H2O, 0.1% Formic Acid (phase B). Separation was achieved with an acetonitrile gradient from 10% to 35% over 360 min, 35% to 100% over 1 min, and 100% B over 5 min at a flow rate of 200 nL/min.

A Thermo ScientificTM Q Exactive™ Plus Orbitrap™ mass spectrometer was used for acquiring the top 10 MS/MS spectra in DDA mode. LC-MS data were analysed using the SEQUEST® HT search engine in Thermo Scientific™ Proteome Discoverer™ 2.2 software considering the modifications: static carbamidomethylation (C), dynamic oxidation (M) and dynamic N-terminus acetylation. Data were searched against the Uniprot *Ustilago maydis* protein database and results were filtered using a 1% protein FDR threshold. Proteins with a -0.5 ≤log2FC≥0.5 and a *p*-value≤0.05 were considered as differentially accumulated in three biological replicates.

#### Sequence Alignment and Phylogenetic Analysis

Phylogenetic analysis of Row family was inferred by using the maximum likelihood method based on the JTT matrix-based model (Jones *et al.*, 1992).

The G-INS progressive method was used to generate *U. maydis* Row family phylogenetic tree. Bootstrap values are higher than 90. Row1 and Row family phylogenetic tree of Ustilaginaceae organism were obtained using G-INS-i (recommended for <200 sequences with global homology with 2 iterative cycles). Bootstrap values are indicated on each branch in Row family phylogenetic tree, and values higher than 90 are indicated with a circle in each branch. Bootstrapping was replicated 1,000 times. Initial trees for the heuristic search were obtained automatically by applying the neighbor-joining algorithm. Taxonomic common tree was developed using NCBI taxonomic browser. Fungi species were annotated as indicated in (Hibbett *et al.*, 2007). BlastP was used to identified Row sequences conservation across fungi. Reciprocal BEST hits blast was used to identify homologs in Ustilaginaceae. Fungi species classification depending on their virulence capacity were developed as previously described (Bauer *et al.*, 2006; Zuo *et al.*, 2019). Trees were visualized and annotated using Interactive Tree of Life (iTOL v6).

#### Predictive analysis tool

To infer protein characteristic, protein sequences were retrieved from the UniProt databases.. InterPro and Expasy-Prosite server was used to identify specific domains (Gasteiger *et al.*, 2003; Blum *et al.*, 2021). We used SignalP (v5.0) (Almagro Armenteros *et al.*, 2019b) to predict the existence of a signal peptide, TMHMM (v2.0c) (Krogh *et al.*, 2001) to predict the existence of transmembrane domains. TargetP (v2.0) (Almagro Armenteros *et al.*, 2019a) was used to predict the location of the protein and ScanProsite (Gattiker *et al.*, 2002; de Castro *et al.*, 2006) to detect the presence of ER retention or mitochondrial motifs and phosphorylated and post-translational modification residues sites. NetNGlyc-1.0 (Julenius, 2007) and NetOGlyc - 4.0 servers (Steentoft *et al.*, 2013) were also used to check glycosylation sites. We scanned EffectorP-fungi 3.0 (Sperschneider & Dodds, 2022) to detect effector proteins and GPI sites were identified using Big-PI (Eisenhaber *et al.*, 2004). For GO annotation we employed Sma3s v2 (Sequence massive annotator using 3 modules) tool (Casimiro-Soriguer *et al.*, 2017). To study protein structures, we used the application of the deep learning-based structure prediction methods AlphaFold (Varadi *et al.*, 2022), visualizing 3D structure in Pymol.

#### Supplementary figures:

***Fig. S1 Δrow1 affects virulence in the genetic backgrounds FB1xFB2, CL13 and SG200.*** Quantification of symptoms in plants infected with Δ*row1* mutant strains at 14 dpi in **(a)** FB1xFB2, **(b)** SG200, and **(c)** CL13 backgrounds. The total number of infected plants is indicated above each column. Two biological replicates were analysed. The Mann–Whitney statistical test was performed for each mutant versus the WT strain (ns, not significant; *$p$-value < 0.05**; $p$-value < 0.5: ***$p$-value < 0.005).

### Figure S1

(a)

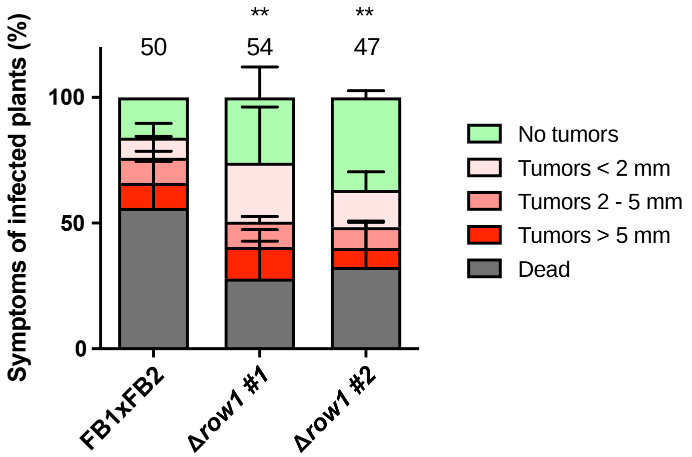


(b)

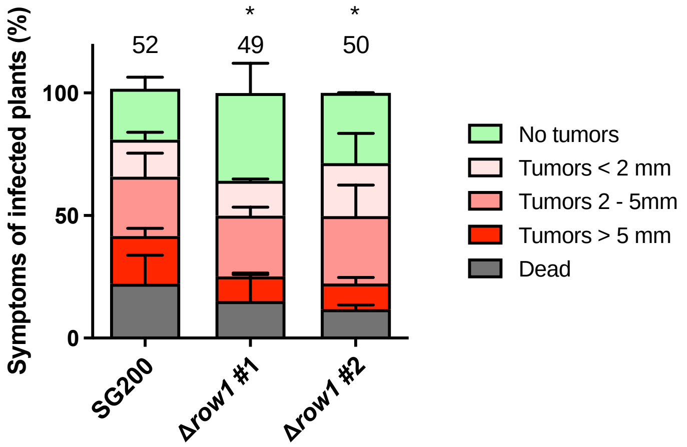


(c)

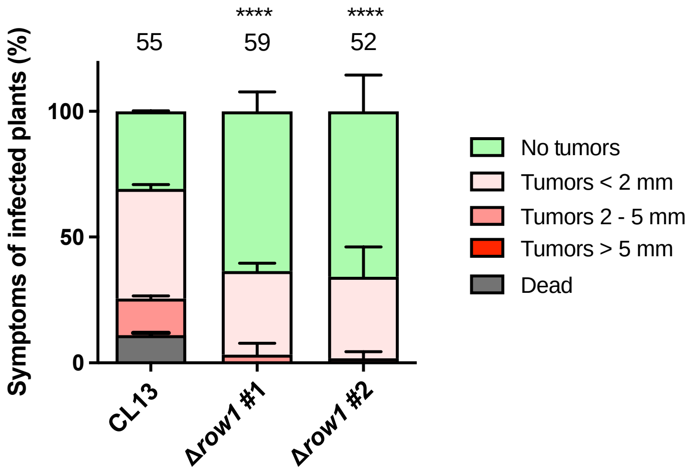

**F*ig. S2 Row1 deletion does not affect growth, cell length or adhesion.*** **(a)** Growth curve of CL13 and Δ*row1* mutant growing in YEPSL media. Error bars represent the standard deviation from three independent replicates. **(b)** Left panel: Cells were visualized by differential interference contrast (DIC) microscopy to analyse cellular morphology. Scale bars represent 10 μm. Right panel: Strains were measured in rich media cultures at exponential phase. Quantification was done for 30 cells from two biological replicates. T-test statistical analysis comparing each mutant versus the WT was performed (ns, no significant). **(c)** Colony adhesion assay of FB1 and Δ*row1* mutant strains. FB1 and Δ*row1* mutants were grown under exponential conditions and applied as droplets onto starch medium plates that were incubated at 28°C for 48 h. Afterward, the surface of the plates was carefully rinsed. The FB1Δ*pmt4* strain was used as a control to assess the impact of reduced fungal cell adhesion to solid surfaces in *U. maydis*. Three biological replicates were performed.

### Figure S2

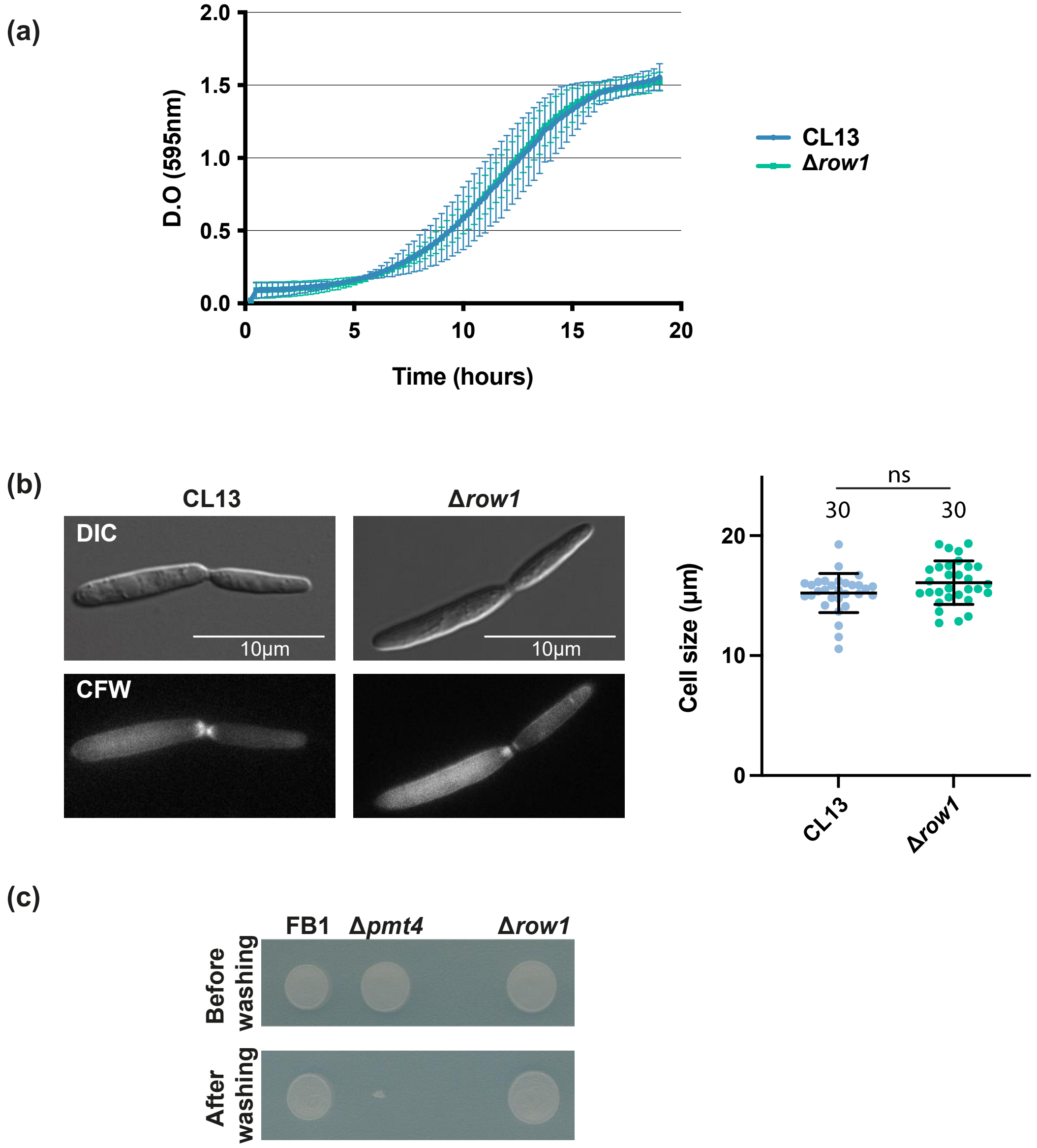

(a)
2.0
1.5
1.0
0.5
0.0
D.O (595nm)
0
5
10
15
20
Time (hours)
CL13
Δrow1
(b)
CL13
Δrow1
DIC
10µm
10µm
CFW
ns
30
30
20
10
0
Cell size (µm)
CL13
Δrow1
(c)
FB1
Δpmt4
Δrow1
Before washing
After washing

***Fig. S3 Δrow1–6 mutants do not present cell growth defects under saline, oxidative, cell wall or reticular stresses.*** A drugs assay was performed in CM plates supplemented with 2% D-glucose, sorbitol 1M and NaCl 0.5M (osmotic stress); $H_2O_2$ 0.75 mM (oxidative stress); calcofluor white (CFW) 10 μg/ml and Congo Red 10 mM (cell wall stress); and Tunicamycin 1 μg/ml (ER stress). The indicated strains were grown under exponential conditions and applied as droplets onto selected plates that were incubated at 28°C for 48 h. A representative picture from three biological replicates performed is shown.

Figure S3

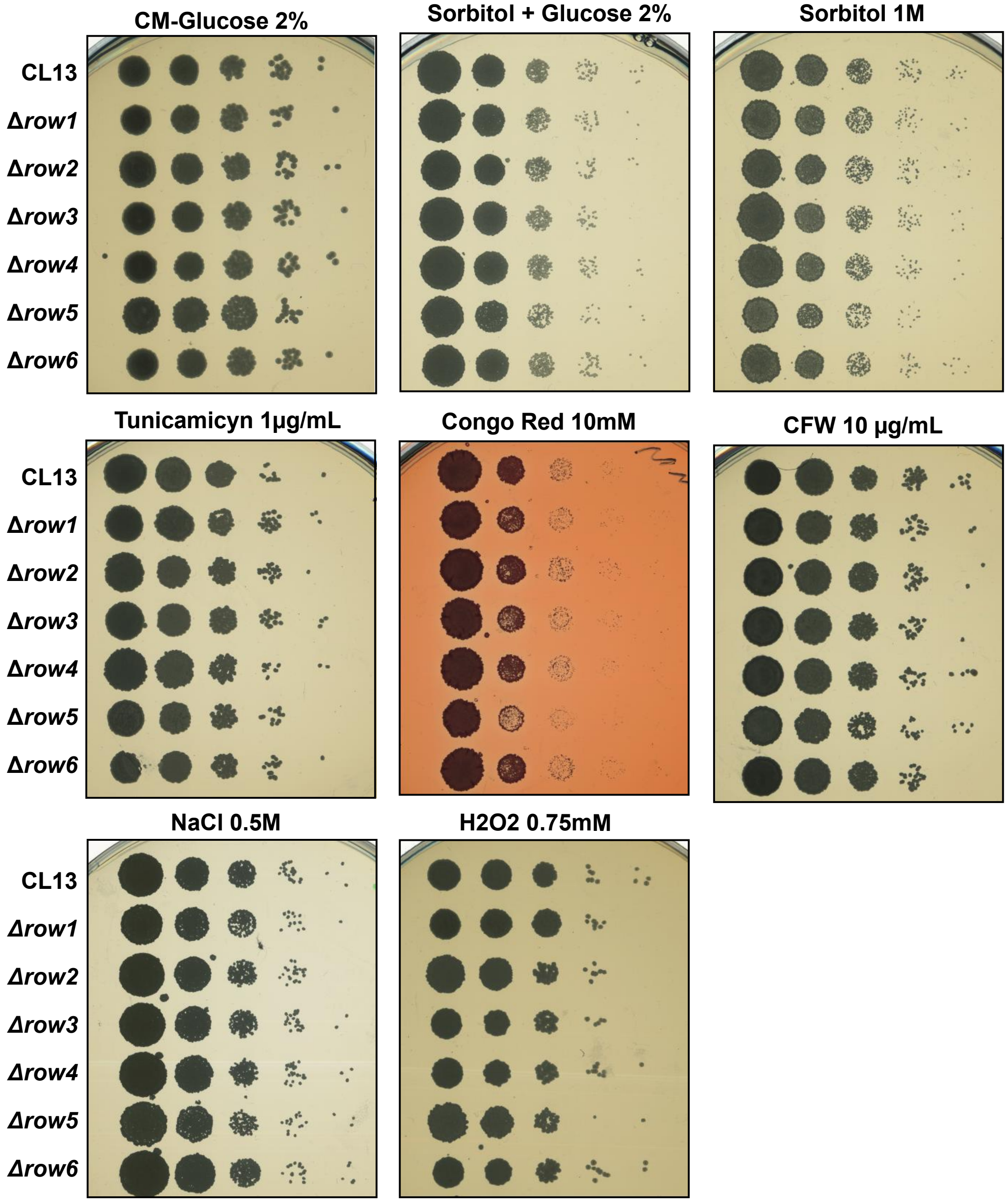
CM-Glucose 2%
Sorbitol + Glucose 2%
Sorbitol 1M
Tunicamicyn 1µg/mL
Congo Red 10mM
CFW 10 µg/mL
NaCl 0.5M
H2O2 0.75mM
CL13
Δrow1
Δrow2
Δrow3
Δrow4
Δrow5
Δrow6

***Fig. S4 Row1 secretion assays.*** **(a)** Secretion of Row1::GFP performed in a colony secretion assay under non-pathogenic and pathogenic conditions using YPDU and PD-charcoal plates, respectively. The SG200 WT strain and SG200 cells expressing cytoplasmic GFP under the control of the constitutive *otef* promoter served as cell lysis controls, and the secreted protein *Cmu1*::GFP was used as a positive control. **(b)** Representation of Row1 architecture tagged with GFP (upper panel). Anti-GFP western blot assay showing cytosolic and secreted protein fraction of AB33 Row1::GFP extracted from cells growing under non-induced conditions and filaments obtained after 5 h of induction (lower panel). Stain-free gel is shown as a loading control.

### Figure S4

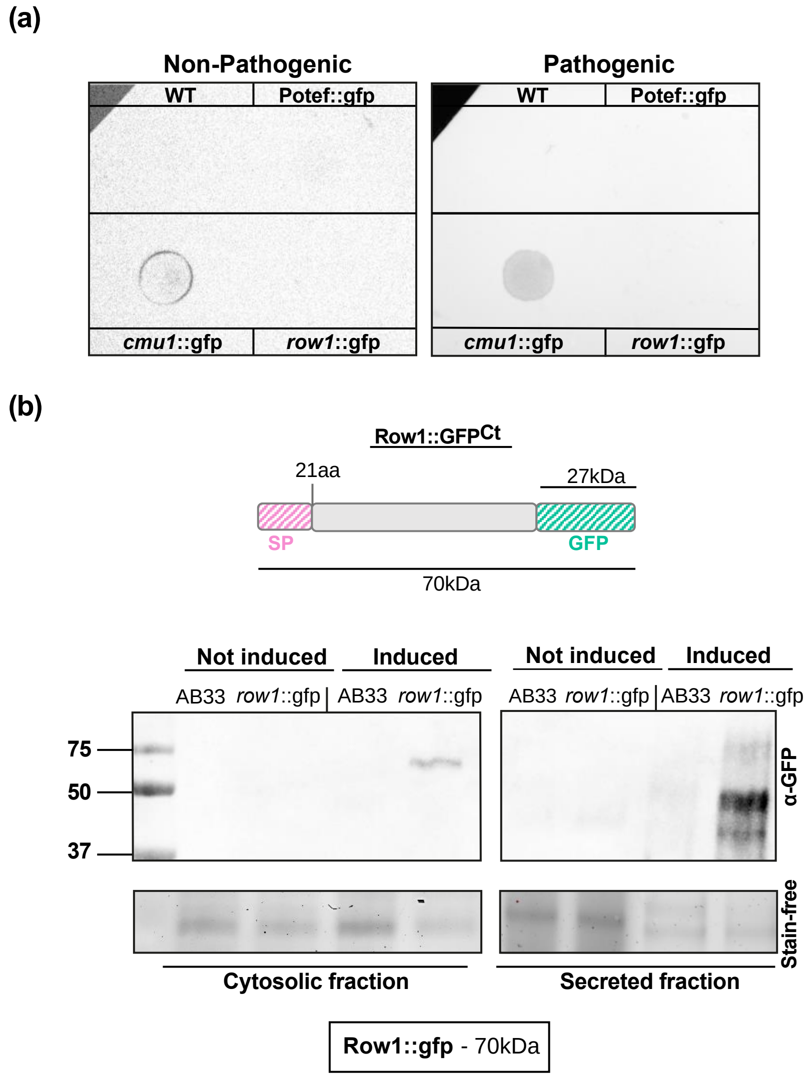

(a)
Non-Pathogenic
Pathogenic
WT
Potef::gfp
cmu1::gfp
row1::gfp
(b)
Row1::GFPCt
21aa
27kDa
SP
GFP
70kDa
Not induced
Induced
AB33
row1::gfp
75
50
37
α-GFP
Stain-free
Cytosolic fraction
Secreted fraction
Row1::gfp - 70kDa

***Fig. S5 The Δrow1 mutant does not exhibit defects in filament length or morphology.*** AB33 and Δ*row1* mutant strains were induced with nitrate for 3 h. Filament length and morphology were visualized by differential interference contrast (DIC) microscopy (upper panel) and quantified (lower panel). Scale bars represent 10 µm. Total number of infected plants is indicated above each column. Error bars represent the standard deviation from three independent replicates. The Student's t-test statistical analysis was performed for mutant versus WT (ns, not significant).

**Figure S5**

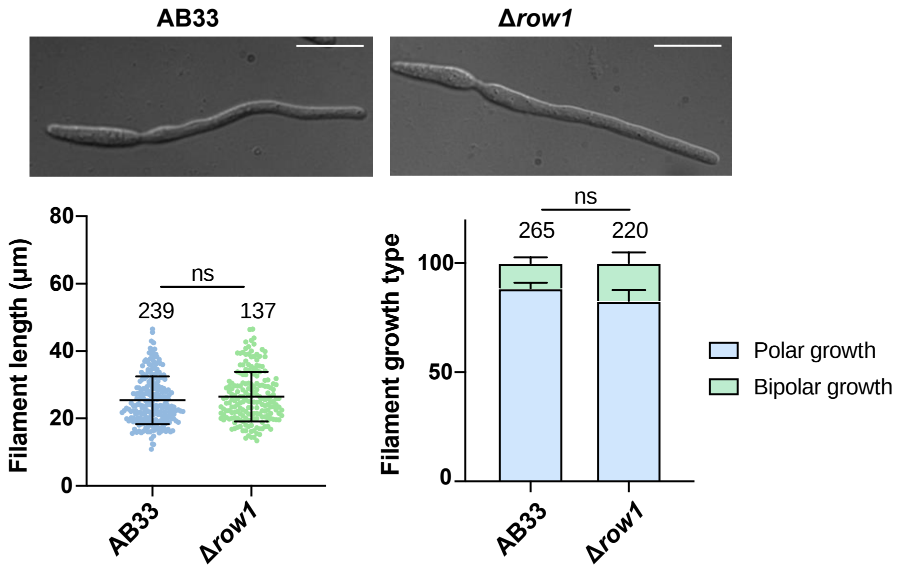

***Fig. S6. Row family sequence alignment.*** Multiple sequence alignment of Row1, Row2, Row3, Row4, Row5 and Row6 using MAFFTv7 and represented by Jalview. Amino acid colours are applied following the Clustal colour scheme: blue (hydrophobic), red (positive charge), magenta (negative charge), green (polar), pink (cysteines), orange (glycines), yellow (prolines), cyan (aromatic), and white (white) (right panel).

**Figure S6**

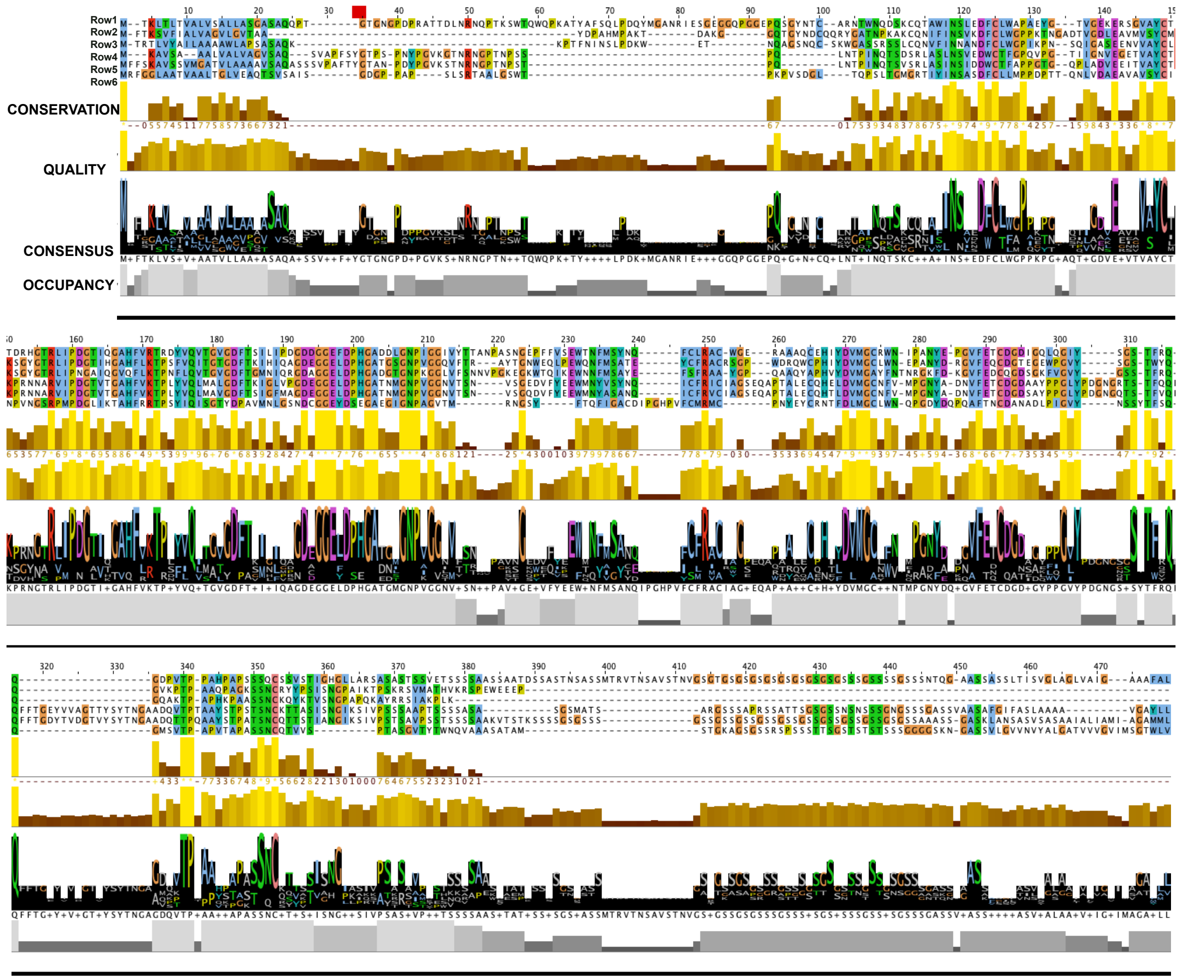

***Fig. S7 Row family structural alignment.*** **(a)** Multiple sequence alignment of Row1, Row2, Row3, Row4 and Row5 focusing on the region spanning approximately amino acids 100–310 obtained using MAFFTv7 and represented by Jalview. Amino acid colours are applied following the Clustal colour scheme: blue (hydrophobic), red (positive charge), magenta (negative charge), green (polar), pink (cysteines), orange (glycines), yellow (prolines), cyan (aromatic), and white (unconserved) (panel right). Superposition of 3D structures corresponding to the indicated region (amino acids 100–310) of each member obtained from AlphaFold PDB and visualized using Pymol (left panel). **(b)** Multiple sequence alignment and structure superposition of the Row family including Row6.

### Figure S7

(a)

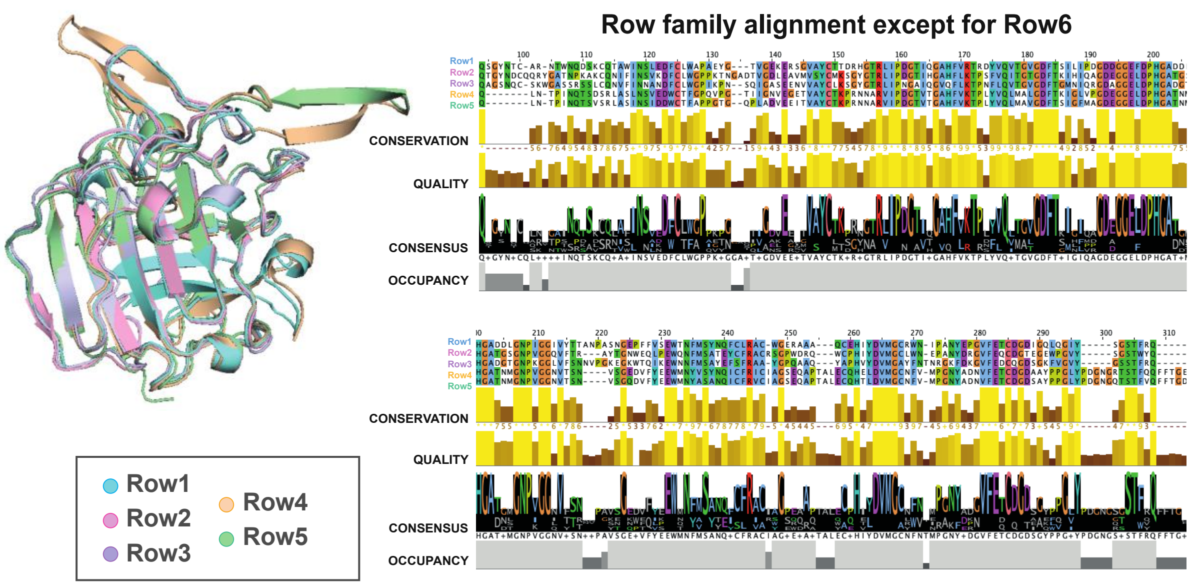
Row family alignment except for Row6
Row1
Row2
Row3
Row4
Row5
CONSERVATION
QUALITY
CONSENSUS
OCCUPANCY
CONSERVATION
QUALITY
CONSENSUS
OCCUPANCY
Row1
Row2
Row3
Row4
Row5

(b)

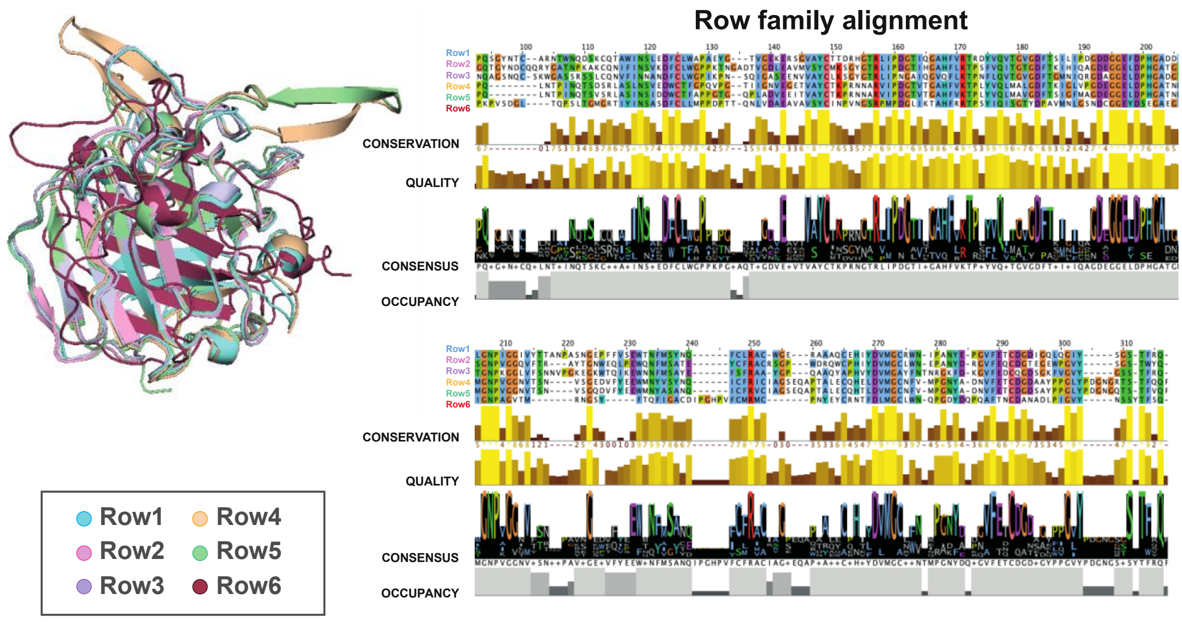
Row family alignment
Row1
Row2
Row3
Row4
Row5
Row6
CONSERVATION
QUALITY
CONSENSUS
OCCUPANCY
CONSERVATION
QUALITY
CONSENSUS
OCCUPANCY
Row1
Row2
Row3
Row4
Row5
Row6

***Fig. S8. The Row family is conserved in Ustilaginales.*** Each member of the *U. maydis* Row family is assigned a specific colour and highlighted: Row1 (pink), Row2 (purple), Row3 (yellow), Row4 (orange), Row5 (blue), and Row6 (green). Homologues of each of these proteins are shown in the same colour, accompanied by the name of the corresponding species and its accession number. BlastP was used to search for homologous sequences in the Ustilaginales order. The alignments were obtained using MAFFT v7. Phylogenetic analysis was inferred by using the maximum likelihood method. Distances are indicated in blue above each branch. The phylogenetic tree was generated using Archaeopteryx.js and edited in iTOL. Bootstrap > 90 is represented with blue circles.

### Figure S8

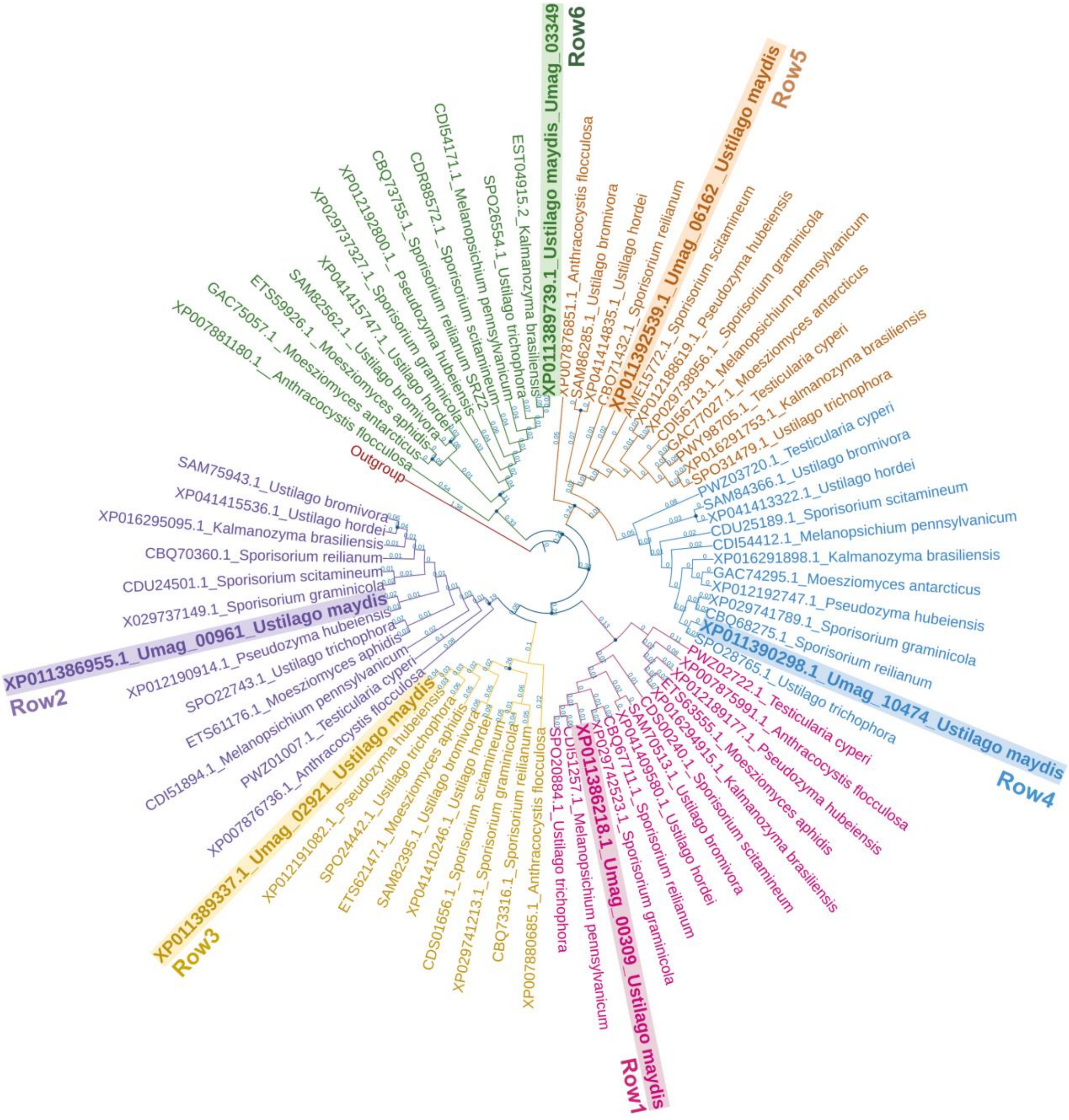
Row1
Row2
Row3
Row4
Row5
Row6
Outgroup
XP011389739.1_Ustilago maydis_Umag_03349
XP011392539.1_Umag_06162_Ustilago maydis
XP011390298.1_Umag_10474_Ustilago maydis
XP011386218.1_Umag_00309_Ustilago maydis
XP011389337.1_Umag_02921_Ustilago maydis
XP011386955.1_Umag_00961_Ustilago maydis
EST04915.2_Kalmanozyma brasiliensis
SPO26554.1_Ustilago trichophora
CDI54171.1_Melanopsichium pennsylvanicum
CDR88572.1_Sporisorium scitamineum
CBQ73755.1_Sporisorium reilianum SRZ2
XP012192800.1_Pseudozyma hubeiensis
XP029737327.1_Sporisorium graminicola
XP041415747.1_Ustilago hordei
SAM82562.1_Ustilago bromivora
ETS59926.1_Moesziomyces aphidis
GAC75057.1_Moesziomyces antarcticus
XP007881180.1_Anthracocystis flocculosa
XP007876851.1_Anthracocystis flocculosa
SAM86285.1_Ustilago bromivora
XP041414835.1_Ustilago hordei
CBQ71432.1_Sporisorium reilianum
AME15772.1_Sporisorium scitamineum
XP012188619.1_Pseudozyma hubeiensis
XP029738956.1_Sporisorium graminicola
CDI56713.1_Melanopsichium pennsylvanicum
GAC77027.1_Moesziomyces antarcticus
PWY98705.1_Testicularia cyperi
XP016291753.1_Kalmanozyma brasiliensis
SPO31479.1_Ustilago trichophora
PWZ03720.1_Testicularia cyperi
SAM84366.1_Ustilago bromivora
XP041413322.1_Ustilago hordei
CDU25189.1_Sporisorium scitamineum
CDI54412.1_Melanopsichium pennsylvanicum
XP016291898.1_Kalmanozyma brasiliensis
GAC74295.1_Moesziomyces antarcticus
XP012192747.1_Pseudozyma hubeiensis
XP029741789.1_Sporisorium graminicola
CBQ68275.1_Sporisorium reilianum
SPO28765.1_Ustilago trichophora
PWZ02722.1_Testicularia cyperi
XP007875991.1_Anthracocystis flocculosa
XP012189177.1_Pseudozyma hubeiensis
ETS63555.1_Moesziomyces aphidis
XP016294915.1_Kalmanozyma brasiliensis
CDS00240.1_Sporisorium scitamineum
SAM70513.1_Ustilago bromivora
XP041409580.1_Ustilago hordei
CBQ67711.1_Sporisorium reilianum
XP029742523.1_Sporisorium graminicola
CDI51257.1_Melanopsichium pennsylvanicum
SPO20884.1_Ustilago trichophora
XP007880685.1_Anthracocystis flocculosa
CBQ73316.1_Sporisorium reilianum
XP029741213.1_Sporisorium graminicola
CDS01656.1_Sporisorium scitamineum
XP041410246.1_Ustilago hordei
SAM82395.1_Ustilago bromivora
ETS62147.1_Moesziomyces aphidis
SPO24442.1_Ustilago trichophora
XP012191082.1_Pseudozyma hubeiensis
XP007876736.1_Anthracocystis flocculosa
PWZ01007.1_Testicularia cyperi
CDI51894.1_Melanopsichium pennsylvanicum
ETS61176.1_Moesziomyces aphidis
SPO22743.1_Ustilago trichophora
XP012190914.1_Pseudozyma hubeiensis
X029737149.1_Sporisorium graminicola
CDU24501.1_Sporisorium scitamineum
CBQ70360.1_Sporisorium reilianum
XP016295095.1_Kalmanozyma brasiliensis
XP041415536.1_Ustilago hordei
SAM75943.1_Ustilago bromivora

t

***Fig. S9. Row family members are involved in U. maydis virulence.*** Quantification of symptoms in plants infected with WT and Δ*row1–6 mutant* strains at 14 dpi. The total number of infected plants is indicated above each column. Two biological replicates were analysed. The Mann–Whitney statistical test was performed for each mutant versus the WT strain (ns, not significant; *p-value < 0.05; **p-value < 0.005; ***$p$-value < 0.0005; and ****$p$-value < 0.0001).

### Figure S9

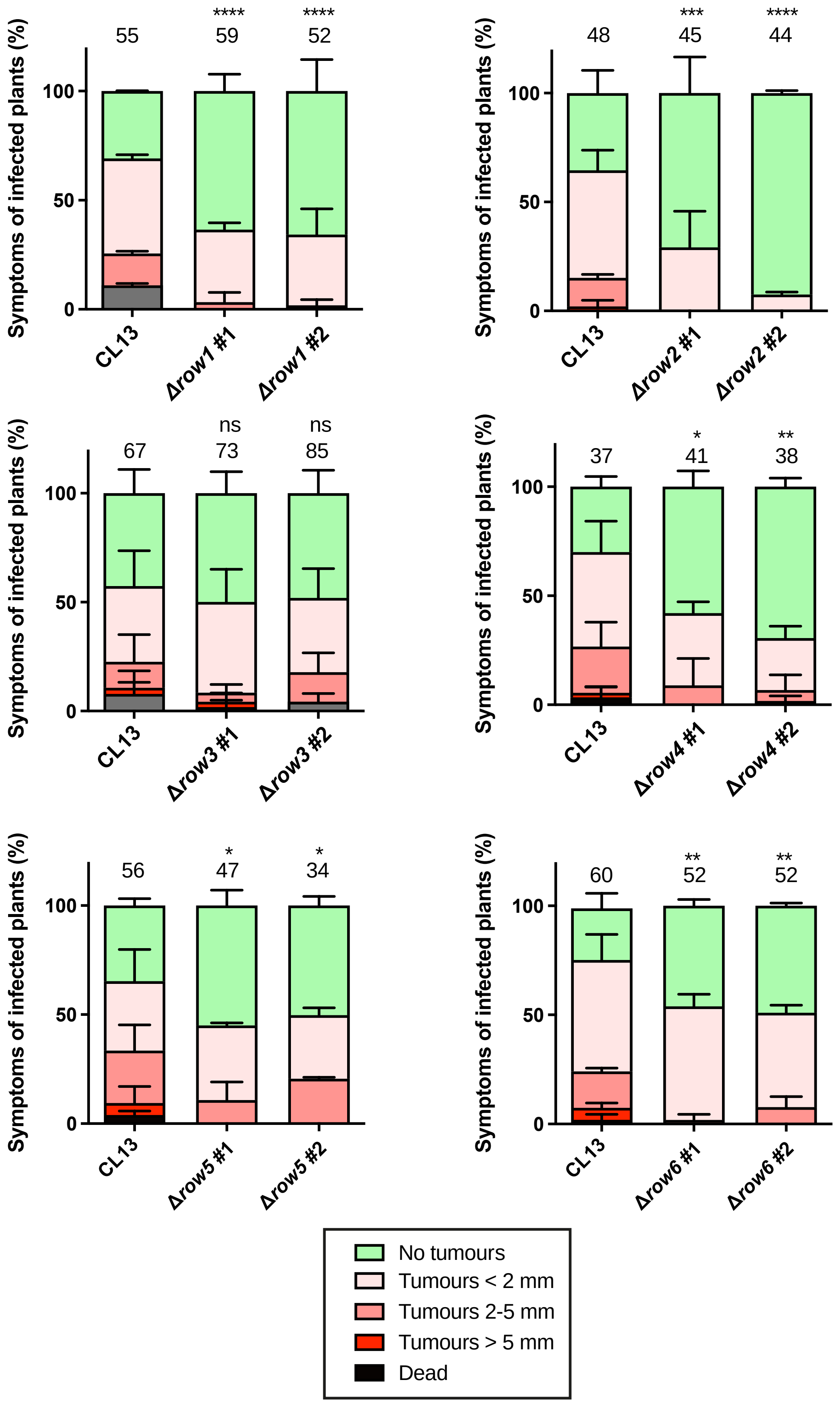
Symptoms of infected plants (%)
0
50
100
55
59
52
****
****
CL13
Δrow1 #1
Δrow1 #2
48
45
44
***
****
CL13
Δrow2 #1
Δrow2 #2
67
73
85
ns
ns
CL13
Δrow3 #1
Δrow3 #2
37
41
38
*
**
CL13
Δrow4 #1
Δrow4 #2
56
47
34
*
*
CL13
Δrow5 #1
Δrow5 #2
60
52
52
**
**
CL13
Δrow6 #1
Δrow6 #2
No tumours
Tumours < 2 mm
Tumours 2-5 mm
Tumours > 5 mm
Dead

***Fig S10. Row family members are differentially expressed during infection.*** Expression levels of *row1*, *row2*, *row3*, *row4*, *row5*, and *row6* represented as normalized counts obtained from high-throughput transcriptomic analysis of *U. maydis* during pathogenesis (Lanver *et al.*, 2018).

**Figure S10**

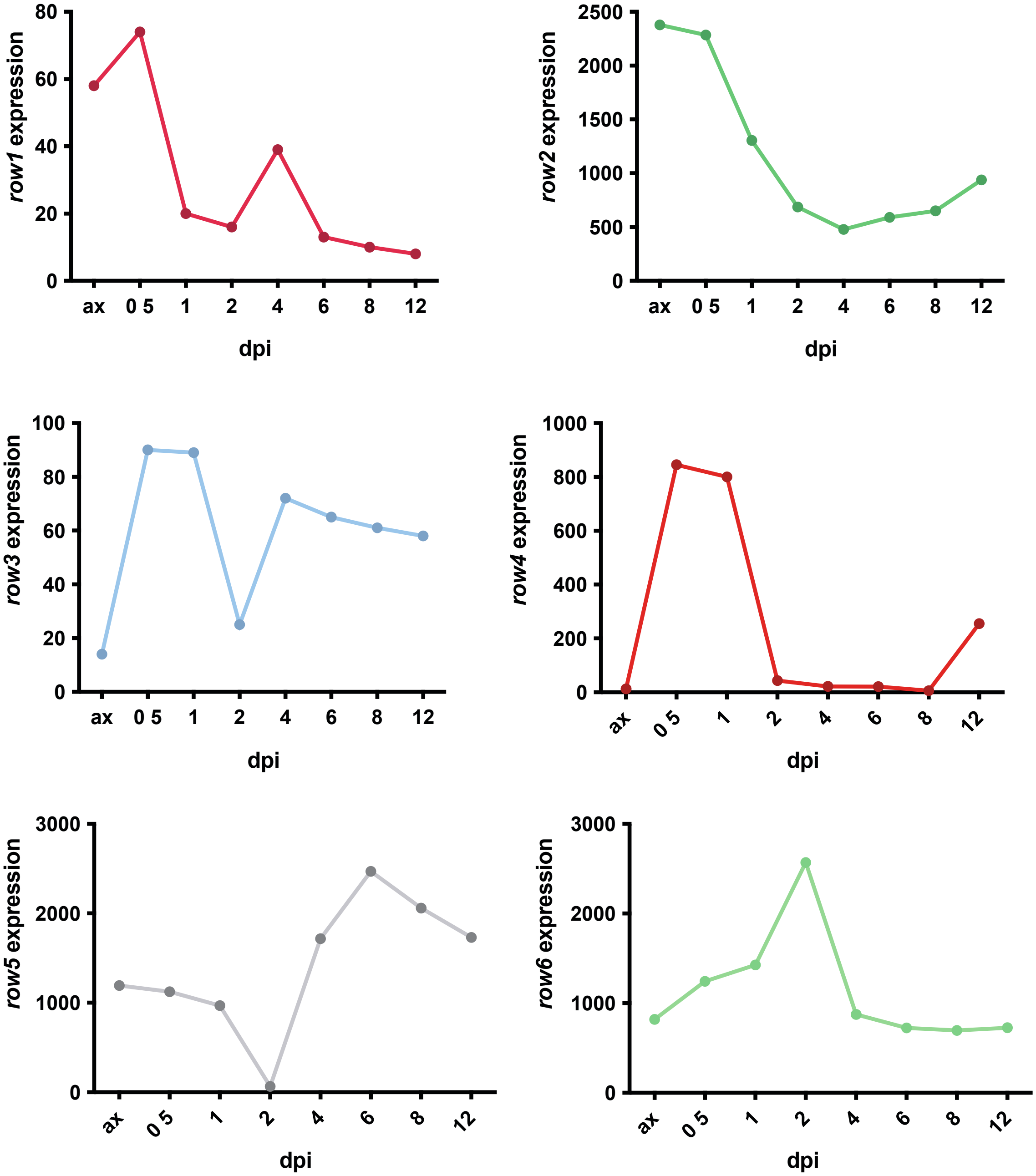

***Fig. S11. Row1 does not have a CBM13 domain, which is conserved in homologues belonging to the Agaricales order.*** Multiple sequence alignment of Row1 of *U. maydis* and homologues of *Amanita muscaria, Mycena chlorophos and Armillaria solidipes* belonging to Agaricales order. Amino acid colours are applied following the Clustal colour scheme: blue (hydrophobic), red (positive charge), magenta (negative charge), green (polar), pink (cysteines), orange (glycines), yellow (prolines), cyan (aromatic), and white (unconserved) (upper panel). Lower panel: Superposition of the 3D structures of Row1 and an *Amanita muscaria* homologue. Row1 is in blue, the *A. muscaria* protein is in pink, and the CBM13 domain of *A. muscaria* is in green.

### Figure S11

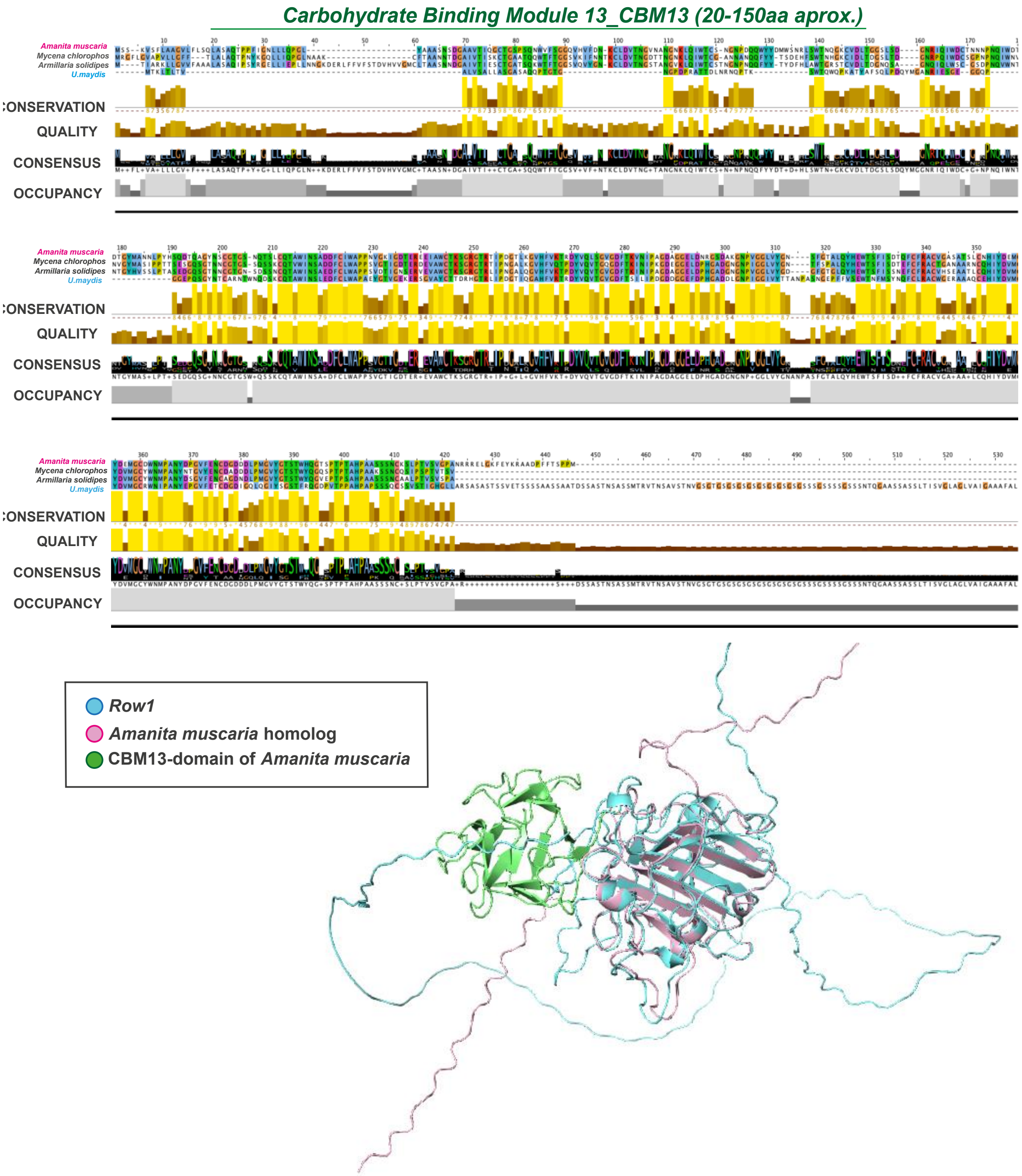

Carbohydrate Binding Module 13_CBM13 (20-150aa aprox.)
Amanita muscaria
Mycena chlorophos
Armillaria solidipes
U.maydis
CONSERVATION
QUALITY
CONSENSUS
OCCUPANCY
Row1
Amanita muscaria homolog
CBM13-domain of Amanita muscaria

**Table S1** Strains used in this study

| Strain | Genotype | Resistance | Source |
|---|---|---|---|
| FB1 | *a1 b1* | - | Banuett and Herskowitz 1989 |
| FB2 | *a2 b2* | - | Banuett and Herskowitz 1989 |
| FB1 Δ*row1* | *a1 b1 Δrow1* | NatR | This work |
| FB2 Δ*row1* | *a2 b2 Δrow1* | NatR | This work |
| CL13 | *a1 bE1 bW2* | | Bolker et al., 1995 |
| CL13 Δ*row1* | *a1 bE1 bW2 Δrow1* | NatR | Marin-Menguiano et al., 2019 |
| AB33 Δ*row1* | *a2 Pnar: bW2 bE1 Δrow1* | NatR | This work |
| CL13 Δ*row1* Prow1:*row1* | *a1 bE1 bW2 Δrow1 ipR[Prow1:row1]ipS* | NatR CbxR | This work |
| CL13 Δ*row1* Prow1:*Sr16611* | *a1 bE1 bW2 Δrow1 ipR[Prow1:Sr16611]ipS* | NatR CbxR | This work |
| CL13 Δ*row1* Prow1:*UHO2_01178* | *a1 bE1 bW2 Δrow1 ipR[Prow1:UHO2_01178]ipS* | NatR CbxR | This work |
| CL13 Δrow2 | *a1 bE1 bW2 Δrow2* | HygR | This work |
| CL13 Δ*row2* Prow2:*row2*::gfp | *a1 bE1 bW2 Δrow2 ipR[Prow2:row2::gfp]ipS* | HygR CxbR | This work |
| CL13 Δ*row3* | *a1 bE1 bW2 Δrow3* | GenR | This work |
| CL13 Δ*row4* | *a1 bE1 bW2 Δrow4* | CxbR | This work |
| CL13 Δ*row5* | *a1 bE1 bW2 Δrow5* | HygR | This work |
| CL13 Δ*row6* | *a1 bE1 bW2 Δrow6* | GenR | This work |
| CL13 Δ*row2*Δ*row1* | *a1 bE1 bW2 Δrow2Δrow1* | NatR HygR | This work |
| CL13 Δ*row1*Δ*row2*Δ*row4* | *a1 bE1 bW2 Δrow2Δrow1Δrow4* | NatR HygR CbxR | This work |
| SG200 | *a1 mfa2 bW2 bE1* | - | Bolker et al., 1995 |
| SG200 Δ*row1* | *a1 mfa2 bW2 bE1 Δrow1* | NatR | Marin-Menguiano et al., 2019 |
| FB1 Δ*pmt4* | *a1 b1 Δpmt4* | HygR | Fernández-Álvarez et al., 2009 |
| SG200 AM1::mCherry | *a1: mfa2 bW2 bE1 ipR[PUMAG_01779:3xegfp]ipS* | CbxR | Mendoza-Mendoza et al., 2009 |
| SG200 AM1::mCherry Δ*row1* | *SG200 AM1::mCherry Δrow1* | CxbR NatR | This work |
| SG200 AM1-Cherry Prow1:*row1*::gfp | *a1: mfa2 bW2 bE1 ipR[PUMAG_01779:3xmCherry Prow1:row1::gfp]ipS* | ? CbxR | This work |
| SG200 Potef:*cmu1*::gfp | *a1 mfa2 bW2 bE1 ipR[Potef:cmu1::gfp]ipS* | CbxR | This work |
| SG200 Potef:*cmu1*::gfp Δrow1 | *a1 mfa2 bW2 bE1 ipR[Potef:cmu1::gfp]ipS Δrow1* | CbxR NatR | This work |
| SG200 Δrow2 Potef:*cmu1*::gfp | *a1: mfa2 bW2 bE1 ipR[Potef:cmu1::gfp]ipS Δrow2* | CbxR HygR | This work |
| AB33 | *a2 Pnar: bW2 bE1* | - | Brachmann et al., 2001 |
| AB33 Prow1:*row1*::gfp | *a2 Pnar: bW2 bE1 ipR[Prow1:row1::gfp]ipS* | CbxR | This work |
| AB33 Prow1:*row1*::gfp mrfp:HDEL | *a2 Pnar: bW2 bE1 ipR[Prow1:row1::gfp Potef:cals::mrfp-HDEL]ipS* | CbxR HygR | This work |

| | | | |
|---|---|---|---|
| AB33 Prow1:*row1*::gfp Potef:yup1::mCherry | *a2 Pnar: bW2 bE1 ipR[Prow1:row1::gfp Potef:yup1::gfp ]ipS* | GenR HygR | This work |
| AB33 Potef:*yup1*::mCherry | *a2 Pnar: bW2 bE1 ipR[Potef:yup1::gfp ]ipS* | GenR | This work |
| AB33 Δ*row1* | *a2 Pnar: bW2 bE1 Δrow1* | NatR | This work |
| AB33 Δ*row1*Δ*row2* | *a2 Pnar: bW2 bE1 Δrow1Δrow2* | NatR HygR | This work |
| SG200 Potef:*row2*::gfp | *a1 mfa2 bW2 bE1 ipR[Potef:row2::*mCherry*]ipS* | CbxR | This work |
| SG200 Potef:*row2*::gfp + Potef:*row1*::mCherry | *a1 mfa2 bW2 bE1 ipR[Potef:row2::mCherry Potef:309::mCherry]ipS* | CbxR GenR | This work |

**Table S2** Plasmids used in this study

| Plasmid | Cloning method | Primer Name | Primer sequence 5'-3' |
|---|---|---|---|
| pJET Δ*row1*:NatR | Standard molecular cloning | CoUm00309-5 | TCGACTAGAATGAATCGGACG |
| | | CoUm00309-3 | GATGGTCGAAACCAAGCAAGC |
| | | Um00309 KO5-1 | ACGACTGGCTGCTAAACAGG |
| | | Um00309 KO5-2 | CACGGCCTGAGTGGCCTGTTGAGAGAGA TGCTTTCCTGC |
| | | Um00309 KO3-1 | GTGGCCATCTAGGCCACTGCGTCGCATGT ATTTCC |
| | | Um00309 KO3-2 | TTCGTGAACCCATCTCCAGC |
| | | Um00309 Int-1 | CGACTTGAATCGCAATCAGC |
| | | Um00309 Int-2 | ACTCGCTAACAAAGAATGGCTCG |
| pJET Δ*row2*:HygR | Standard molecular cloning | Co5_Um00961 | GAACGACAAGCCTTTTTG |
| | | Co3_Um00961 | GAGCATAACTTACCTCCT |
| | | Ko5-1 Um00961 | TTCAGCTGACACCAGTCA |
| | | Ko5-2 Um00961 | CACGGCCTGAGTGGCCCAGTCGATTGGA ATTGT |
| | | KO3-1_Um00961 | GTGGCCATCTAGGCCGTCGCTCCTTCCAA AGTC |
| | | KO3-2_Um00961 | GTCCTTGATCGAAGAGAA |
| | | INT5_II_Umag_00961 | CCAAGACCAATGGCGCCG |
| | | INT3_II_Umag_00961 | CACCGACGGGGTTACCAG |
| pBSK Δ*row3*:GenR | NEBuilder® HiFi DNA Assembly | Ko5' Umag_02921_fwd | cgaattcctgcagcccggggTTGAAAAATTCTCTCA TCTAATCAATCAC |
| | | Ko5' Umag_02921_rev | acgccatggtGGAGGTCTTCTACCGACAC |
| | | GenR_fwd (2921) | gaagacctccACCATGGCGTGACAATTG |
| | | GenR_rev (2921) | cggcccgagtTATTAATGCGGCCGCACTC |
| | | Ko3' Umag_02921_fwd | cgcattaataACTCGGGCCGCTCGTAGC |
| | | Ko3' Umag_02921_rev | cggccgctctagaactagtgTTGGATTGGTGCCGAG AACTTTCC |
| | | Co5' Umag_2921_fwd | TTGGCGGATTTGATCAGG |
| | | Co3' Umag_02921_rev | CGAGAAATCATGAATGGA |
| | | INT5' Umag_2921 | ACGATTTCTGCCTATGGG |
| | | INT3' Umag_2921 | TAGGCGCCCATGACATCG |

| | | | |
|---|---|---|---|
| | | Ko5-1_Umag02921_PCR | TTGAAAAATTCTCTCATC |
| | | Ko3-2_Umag02921_PCR | TTGGATTGGTGCCGAGAA |
| pBSK Δ*row5*::HygR | NEBuilder® HiFi DNA Assembly | Ko5' Umag_06162_fwd | cgaattcctgcagcccggggTATTGTGAATAGGATTAATTCTAGGC |
| | | Ko5' Umag_06162_rev | cgttcggccaAGGAGAGGAGGAGAAAGAG |
| | | HygR_fwd_6162 | ctcctctcctTGGCCGAACGTGGTAACTAC |
| | | HygR_rev_6162 | aaacgttgtcCTATTCCTTTGCCCTCGG |
| | | Ko3'-1 Umag_06162_fwd | aaaggaatagGACAACGTTTCGCCCTCC |
| | | Ko3'-1 Umag_06162_rev | cggccgctctagaactagtgTACGCTCGACTTTCTCTACG |
| | | Co5’ Umag_06162 | ATTTCACTCGTGACGGAC |
| | | Co3’ Umag_06162 | GATTTTGCACGATCCAAG |
| | | INT5’ Umag_06162 | TTTCAGCCCGACGGAACCG |
| | | INT3’ Umag_06162 | CGGTAGGAGTGACCTGGT |
| | | Ko5-1 Umag_06162_PCR | TATTGTGAATAGGATTAA |
| | | Ko3-2_Umag06162_PCR_II | tacgctcgactttctcta |
| pBSK Δ*row4*:CbxR | NEBuilder® HiFi DNA Assembly | Ko5' Umag_10474_fwd | cgaattcctgcagcccggggGTGGTGCGTGGTGCGTGG |
| | | Ko5' Umag_10474_rev | gctcgatattGATGGTGGAGAGAAGCAAGACAAGG |
| | | CbxR_fwd (10474) | ctccaccatcAATATCGAGCACGTTGATGG |
| | | CbxR_rev (10474) | tcgtctttgtAATGGGGATCTTCGCTCAAC |
| | | Ko3' Umag_10474_fwd | gatccccattACAAAGACGATGACCAAG |
| | | Ko3' Umag_10474_rev | tcgtctttgtAATGGGGATCTTCGCTCAAC |
| | | Co5’ Umag_10474 | CCAAAACCAGTTGACAGA |
| | | Co3’ Umag_10474 | CTAGATCGGTGCGCGAAA |
| | | INT5’ Umag_10474 | GCGTCATCCCCGACGGTA |
| | | INT3’ Umag_10474 | CGAAGTCTGACCGTTGCC |
| | | Ko5-1 Umag_10474_PCR | GTGGTGCGTGGTGCGTGG |
| | | Ko3-2 Umag_10474_PCR | CTTCGAATGCCAAGTTGA |
| pJET Δ*row6*::GenR | Standard molecular cloning | Co5-Um03349 | TGTATTGCTGAGCCTTGT |
| | | KO5-1_Um03349 | GAAGGCCACAACTCGTGA |
| | | KO5-2_Um03349 | CACGGCCTGAGTGGCCGACGGTGATGAACACGAC |
| | | INT5_Um03349 | AGGCGGTCGCTGTGTCAT |
| | | INT3_Um03349 | TAACAGGCGCAGGAGTGA |
| | | KO3-1_Um03349 | GTGGCCATCTAGGCCTTCTGTGTTCGTGGTTC |
| | | KO3-2_Um03349 | ATGATTATTCGTTCTTCG |
| | | Co3-Um03349 | TGCCGAGTTCAAAGTTGT |

| | | | |
|---|---|---|---|
| p123 Prow1:*row1* | Standard molecular cloning | Fwd-UMAG_00309 Promoter (PuvII) | TCTCGAGGGCATCGACAA |
| | | Rev-UMAG_00309 (NotI) pOTEF not GFP | AGCgcggccgcTTAGAGAGCAAACGCCGCA |
| p123 Prow1:*row1*::gfp | Standard molecular cloning | Fwd-UMAG_00309 Promoter (PuvII) | TCTCGAGGGCATCGACAA |
| | | Rev-UMAG_00309 (NcoI) | AGCccatggATGTGCAATCTTGTCGGA |
| p123 Prow1::gfp | Standard molecular cloning | Fwd-UMAG_00309 Promoter (PuvII) | TCTCGAGGGCATCGACAA |
| | | Prom309_NcoI | AGCCCATGGGGTGTTGAGAGAGATGCT |
| p123 Prow1:*UHO2_01178* | NEBuilder® HiFi DNA Assembly | Fwd_UHO2_01178_NEB | aagcatctctctcaacacccATGTCCAGCATCAAGACTGCAG |
| | | Rev_UHO2_01178_NEB | tgaacgatctgcagccgggcTCAAACCGGCGGCACAGA |
| p123 Prow1:*Sr16611* | NEBuilder® HiFi DNA Assembly | Fwd_Sr11661_NEB | aagcatctctctcaacacccATGACCAAAATCGCGCTC |
| | | Rev_Sr116611_NEB | tgaacgatctgcagccgggcTCAAAGGGCAAGCACAGC |
| p123 Prow2:*row2* | Standard molecular cloning | Fwd-P961_KpnI | TATggtaccTGTTGCTCGGTGGATATG |
| | | Rev-P961_NotI | AGCgcggccgcTCAAGGCTCCTCCTCCCA |
| p123 Potef:*yup1*::mCherry GenR | NEBuilder® HiFi DNA Assembly | Yup1-Fwd (NEB) | aacatcatccacgggatcccATGGCACAAACACAGCCAC |
| | | Yup1-Rev (NEB) | tgctcaccatAAACTTTCTTCCTATCCCAGC |
| | | mCherry-GenR_fwd (yup1) | aagaaagtttATGGTGAGCAAGGGCGAG |
| | | mCherry-GenR_rev (yup1) | tgaacgatctgcagccgggcGCAAATTAAAGCCTTCGAGCG |
| p123 Potef:*row2*::gfp | NEBuilder® HiFi DNA Assembly | Umag_00961_XmaI_NEB | AACATCATCCACGGGATCCCCCGGGATGTTCACCAAGTCTGTTTTCATCG |
| | | Umag_00961_NcoI_NEB | AGCTCCTCGCCCTTGCTCACCATGGAAGGCTCCTCCTCCCACTC |
| p123 Potef:*row1*::mCherry GenR | NEBuilder® HiFi DNA Assembly | Umag_00309_Fwd_NEB | aacatcatccacgggatcccATGACCAAACTCACGCTCACC |
| | | Umag_00309_Rev_NEB | tgctcaccatGAGAGCAAACGCCGCAGC |
| | | mCherry-GenR_NEB_Fwd (309) | gtttgctctcATGGTGAGCAAGGGCGAG |
| | | mCherry-GenR_NEB_Rev (309) | tgaacgatctgcagccgggcGGCCACTCAGGCCTATTAATG |

**Table S3** Accession numbers

| Protein name | Accession numer (NCBI) | UMAG |
|---|---|---|
| Row1 | XP_011386218.1 | UMAG_00309 |
| Row2 | XP_011386955.1 | UMAG_00961 |
| Row3 | XP_011389337.1 | UMAG_02921 |
| Row4 | XP_011390298.1 | UMAG_10474 |
| Row5 | XP_011392539.1 | UMAG_06162 |
| Row6 | XP_011389739.1 | UMAG_03349 |
| Pmt4 | XP_011392118.1 | UMAG_05433 |
| Cmu1 | XP_011391476.1 | UMAG_05731 |

**Table S5** Protein homologues of Row1 in *U. maydis*

| Protein Name | Coverage (%) | e-value | Identity (%) | Accession | Organism | Coverage (%) | e-value |
|---|---|---|---|---|---|---|---|
| UMAG_00309 | 100 | 0.0 | 78.69% | CBQ67711.1 | *Sporisorium reilianum* | 100 | 0.0 |
| UMAG_00961 | 100 | 0.0 | 80.28% | CDS00240.1 | *Sporisorium scitamineum* | 100 | 0.0 |
| UMAG_02921 | 100 | 0.0 | 80.28% | SPO19971.1 | *Sporisorium trichophora* | 100 | 0.0 |
| UMAG_06162 | 100 | 0.0 | 78.17% | XP_029742523 | *Sporisorium graminicola* | 100 | 0.0 |
| UMAG_10474 | 100 | 0.0 | 76.89% | XP_012189177.1 | *Pseudozyma hubeiensis* | 100 | 0.0 |
| UMAG_3349 | 100 | 0.0 | 76.94% | XP_041409580.1 | *Ustilago hordei* | 100 | 0.0 |

**Table S7** Protein homologues of Row1 in Ustilaginaceae.

| Organism | Coverage (%) | e-value | Identity (%) | Accession |
|---|---|---|---|---|
| *Sporisorium reilianum* | 100 | 0.0 | 78.69% | CBQ67711.1 |
| *Sporisorium scitamineum* | 100 | 0.0 | 80.28% | CDS00240.1 |
| *Sporisorium trichophora* | 100 | 0.0 | 80.28% | SPO19971.1 |
| *Sporisorium graminicola* | 100 | 0.0 | 78.17% | XP_029742523 |
| *Pseudozyma hubeiensis* | 100 | 0.0 | 76.89% | XP_012189177.1 |
| *Ustilago hordei* | 100 | 0.0 | 76.94% | XP_041409580.1 |
| *Ustilago bromivora* | 80 | 0.0 | 83.76% | SAM70513.1 |
| *Melanopsichium pennsylvanicum* | 100 | 0.0 | 75.53% | CDI51257.1 |
| *Moesziomyces antarticus* | 54 | 8E-74 | 53.14% | ETS61176.1 |
| *Moesziomyces aphidis* | 73 | 0.0 | 86.90% | ETS63555.1 |
| *Kalmanozyma brasiliensis* | 100 | 0.0 | 73.65% | XP_016294915.1 |
| *Anthracocystis flocculosa* | 66 | 5E-177 | 83,45% | XP_007875991.1 |
| *Testicularia cyperi* | 73 | 3E-179 | 77.88% | PWZ02722.1 |
| *Violaceomyces palustris* | 73 | 5E-81 | 54.98% | PWN48520.1 |

**Table S8** Row family conservation in Ustilaginaceae.

| ORGANISM | Row members conservation | | | | | |
|---|---|---|---|---|---|---|
| | **Row1** | **Row2** | **Row3** | **Row4** | **Row5** | **Row6** |
| *Ustilago maydis* | | | | | | |
| *Sporisorium scitamineum* | | | | | | |
| *Sporisorium reilianum SRZ2* | | | | | | |
| *Sporisorium graminicola* | | | | | | |
| *Ustilago trichophora* | | | | | | |
| *Ustilaho hordei* | | | | | | |
| *Ustilago bromivora* | | | | | | |
| *Pseudozyma hubeiensis SY62* | | | | | | |
| *Moesziomyces aphidis DSM 70725* | | | | x | x | |
| *Moesziomyces antarcticus T-34* | | | | | | |
| *Melanopsichium pennsylvanicum 4* | | | x | | | |
| *Testicularia cyperi* | | | x | | | x |
| *Anthracocystis flocculosa PF-1* | | | | | | |
| *Kalmanozyma brasiliensis GHG001* | | | x | | | x |

***The green boxes indicate the conservation of each protein in the specified species. The red boxes with a cross indicate their absence in those species.*

**Table S9** Protein homologues of Row family members in Ustilaginaceae.

| Organism | Coverage (%) | e-value | Identity (%) | Acession number |
|---|---|---|---|---|
| ***Ustilago maydis*_Row1** | **100%** | **0.0** | **100.00%** | **XP_011386218.1** |
| *Sporisorium scitamineum* | 100% | 0.0 | 80.28% | CDS00240.1 |
| *Ustilago trichophora* | 100% | 0.0 | 79.81% | SPO20884.1 |
| *Pseudozyma hubeiensis SY62* | 100% | 0.0 | 76.89% | XP_012189177.1 |
| *Ustilago hordei* | 100% | 0.0 | 76.94% | XP_041409580.1 |
| *Sporisorium reilianum SRZ2* | 100% | 0.0 | 78.69% | CBQ67711.1 |
| *Moesziomyces antarcticus T-34* | 73% | 0.0 | 86,90% | GAC73538.1 |
| *Ustilago bromivora* | 80% | 0.0 | 83.76% | SAM70513.1 |
| *Sporisorium graminicola* | 100% | 0.0 | 78.17% | XP_029742523.1 |
| *Melanopsichium pennsylvanicum 4* | 100% | 0.0 | 75.53% | CDI51257.1 |
| *Kalmanozyma brasiliensis GHG001* | 100% | 0.0 | 73.65% | XP_016294915.1 |
| *Moesziomyces aphidis DSM 70725* | 73% | 0.0 | 86.90% | ETS63555.1 |
| *Testicularia cyperi* | 73% | 0.0 | 79.17% | PWZ02722.1 |
| *Anthracocystis flocculosa PF-1* | 66% | 5E-177 | 83.45% | XP_007875991.1 |
| ***Ustilago maydis*_Row2** | **100%** | **0.0** | **100.00%** | **XP_011386955.1** |
| *Pseudozyma hubeiensis SY62* | 100% | 0.0 | 96.55% | XP_012190914.1 |
| *Sporisorium scitamineum* | 99% | 0.0 | 94.46% | CDU24501.1 |
| *Sporisorium reilianum SRZ2* | 100% | 0.0 | 94.14% | CBQ70360.1 |
| *Sporisorium graminicola* | 100% | 0.0 | 91.72% | XP_029737149.1 |
| *Ustilago trichophora* | 100% | 0.0 | 91.38% | SPO22743.1 |
| *Melanopsichium pennsylvanicum 4* | 100% | 0.0 | 91.38% | CDI51894.1 |
| *Kalmanozyma brasiliensis GHG001* | 100% | 0.0 | 90.69% | XP_016295095.1 |
| *Moesziomyces antarcticus T-34* | 99%% | 0.0 | 87,89% | GAC76095.1 |
| *Moesziomyces aphidis DSM 70725* | 99% | 0.0 | 87.89% | ETS61176.1 |
| *Ustilago hordei* | 100% | 0.0 | 88.28% | XP_041415536.1 |

| | | | | |
|---|---|---|---|---|
| *Ustilago bromivora* | 100% | 0.0 | 87.24% | SAM75943.1 |
| *Testicularia cyperi* | 99% | 3E-179 | 80.28% | PWZ01007.1 |
| *Anthracocystis flocculosa PF-1* | 99% | 3E-176 | 78.62% | XP_007876736.1 |
| ***Ustilago maydis*_Row3** | **100%** | **0.0** | **100.00%** | **XP_011389337.1** |
| *Pseudozyma hubeiensis SY62* | 92% | 8E-180 | 89.35% | XP_012191082.1 |
| *Moesziomyces antarcticus T-34* | 98% | 5E-167 | 80.78% | GAC76095.1 |
| *Sporisorium graminicola* | 100% | 1E-177 | 85.96% | XP_029741213.1 |
| *Ustilago trichophora* | 100% | 2E-177 | 82.46% | SPO24442.1 |
| *Sporisorium reilianum SRZ2* | 92% | 2E-173 | 85.93% | CBQ73316.1 |
| *Moesziomyces aphidis DSM 70725* | 98% | 8E-166 | 80.78% | ETS62147.1 |
| *Ustilago bromivora* | 91% | 6E-130 | 69.81% | SAM82395.1 |
| *Anthracocystis flocculosa PF-1* | 95% | 4E-108 | 59.78% | XP_007880685.1 |
| *Sporisorium scitamineum* | 92%% | 8E-176 | 87.07% | CDS01656.1 |
| *Anthracocystis flocculosa PF-1* | 95%% | 5E-109 | 59.78% | XP_007880685.1 |
| *Ustilago hordei* | 78% | 2E-98 | 66.07% | XP_041410246.1 |
| ***Ustilago maydis*_Row4** | **100%** | **0.0** | **100.00%** | **XP_011390298.1** |
| *Pseudozyma hubeiensis SY62* | 75% | 0.0 | 91.26% | XP_012192747.1 |
| *Sporisorium scitamineum* | 76% | 0.0 | 88.39% | CDU25189.1 |
| *Ustilago trichophora* | 75% | 0.0 | 88.42% | SPO28765.1 |
| *Sporisorium graminicola* | 76% | 0.0 | 88.71% | XP_029741789.1 |
| *Sporisorium reilianum SRZ2* | 76% | 0.0 | 88.06% | CBQ68275.1 |
| *Moesziomyces antarcticus T-34* | 75% | 0.0 | 88.35% | GAC74295.1 |
| *Ustilago hordei* | 75% | 0.0 | 89.00% | XP_041413322.1 |
| *Kalmanozyma brasiliensis GHG001* | 75% | 0.0 | 87.06% | XP_016291898.1 |
| *Ustilago bromivora* | 75% | 0.0 | 89.00% | SAM84366.1 |
| *Testicularia cyperi* | 70% | 1E-173 | 77.97 | PWZ03720.1 |
| *Anthracocystis flocculosa PF-1* | 74% | 6E-172 | 74.75% | XP_007882513.1 |
| *Melanopsichium pennsylvanicum 4* | 75% | 0.0 | 83.50% | CDI54412.1 |
| ***Ustilago maydis*_Row5** | **100%** | **0.0** | **100.00%** | **XP_011392539.1** |
| *Moesziomyces antarcticus T-34* | 75% | 0.0 | 93.01% | GAC77027.1 |
| *Ustilago trichophora* | 74% | 0.0 | 92.25% | SPO31479.1 |
| *Sporisorium reilianum SRZ2* | 95% | 0.0 | 82.87% | CBQ71432.1 |
| *Pseudozyma hubeiensis SY62* | 74% | 0.0 | 91.87% | XP_012188619.1 |
| *Melanopsichium pennsylvanicum 4* | 96% | 0.0 | 80.65% | CDI56713.1 |
| *Testicularia cyperi* | 96% | 0.0 | 79.56% | PWY98705.1 |
| *Sporisorium scitamineum* | 73% | 0.0 | 91.79% | AME15772.1 |
| *Kalmanozyma brasiliensis GHG001* | 75% | 0.0 | 88.19% | XP_016291753. |
| *Anthracocystis flocculosa PF-1* | 100% | 0.0 | 72.35% | XP_007882513.1 |
| *Sporisorium graminicola* | 74% | 0.0 | 89.08% | XP_029738956.1 |
| *Ustilago hordei* | 96% | 0.0 | 78.26% | XP_041414835.1 |
| *Ustilago bromivora* | 74% | 0.0 | 87.37% | SAM86285.1 |
| ***Ustilago maydis*_Row6** | **100%** | **0.0** | **100.00%** | **XP_011389739.1** |
| *Pseudozyma hubeiensis SY62* | 95%% | 0.0 | 79.04% | XP_012192800.1 |
| *Ustilago trichophora* | 100% | 0.0 | 78.57% | SPO26554.1 |
| *Moesziomyces aphidis DSM 70725* | 99% | 0.0 | 75.93% | ETS59926.1 |
| *Ustilago bromivora* | 100% | 0.0 | 75.07% | SAM82562.1 |
| *Moesziomyces antarcticus T-34* | 99%% | 0.0 | 75.93% | GAC75057.1 |
| *Ustilago hordei* | 100% | 0.0 | 75.64% | XP_041415747.1 |
| *Melanopsichium pennsylvanicum 4* | 100% | 0.0 | 74.29% | CDI54171.1 |
| *Sporisorium scitamineum* | 78% | 2E-179 | 85.51% | CDR88572.1 |

| | | | | |
|---|---|---|---|---|
| *Sporisorium graminicola* | 78% | 8E-174 | 84.06% | XP_029737327.1 |
| *Kalmanozyma brasiliensis GHG001* | 95% | 4E-171 | 73.00% | EST04915.2 |
| *Sporisorium reilianum SRZ2* | 79% | 1E-164 | 82.14% | CBQ73755.1 |
| *Anthracocystis flocculosa PF-1* | 77% | 1E-86 | 48.72% | XP_007881180.1 |

**Table S11** Homologues of Row1 in *Cryptococcus neoformans*.

| Organism | Name | Accession number | U.maydis homolog | Max Score | Total Score | Query Cover | E-value | Identity | Length |
|---|---|---|---|---|---|---|---|---|---|
| Cryptococcus neoformans var. grubii H99" | CNAG_00776 Immunoreactive mannoprotein MP88 | XP_012046953.1 | Umag_06162 | 333 | 333 | 70% | 5,00E-113 | 58.80% | 381 |
| | | | Umag_10474 | 332 | 332 | 73% | 4E-112 | 57.14% | 407 |
| | CNAG_6000 Glycoprotein | XP_012052961.1 | Umag_06162 | 300 | 300 | 55% | 2,00E-98 | 54.74% | 381 |
| | | | umag_10474 | 274 | 274 | 55% | 3,00E-88 | 51.09% | 407 |
| | CNAG_05312 Hypothetical protein | XP_012048592.1 | Umag_00961 | 254 | 254 | 57% | 2E-82 | 49.63% | 290 |
| | | | Umag_00309 | 224 | 224 | 50% | 4E-69 | 50.00% | 424 |
| | CNAG_07760 Hypothetical protein | XP_012051878.1 | Umag_00961 | 127 | 127 | 58% | 1E-35 | 39.44% | 290 |

#### Supporting information references


**Almagro Armenteros JJ, Salvatore M, Emanuelsson O, Winther O, Heijne von G, Elofsson A, Nielsen H**. **2019a**. Detecting sequence signals in targeting peptides using deep learning. *Life science alliance* **2**: 1–14.

**Almagro Armenteros JJ, Tsirigos KD, Sønderby CK, Petersen TN, Winther O, Brunak S, Heijne von G, Nielsen H**. **2019b**. SignalP 5.0 improves signal peptide predictions using deep neural networks. *Nature Biotechnology* **37**: 420–423.

**Bauer R, Begerow D, Sampaio JP, Weiβ M, Oberwinkler F**. **2006**. The simple-septate basidiomycetes: a synopsis. *Mycological Progress* **5**: 41–66.

**Blum M, Chang H-Y, Chuguransky S, Grego T, Kandasaamy S, Mitchell A, Nuka G, Paysan-Lafosse T, Qureshi M, Raj S, *et al.* 2021**. The InterPro protein families and domains database: 20 years on. *Nucleic acids research* **49**: D344–D354.

**Bösch K, Frantzeskakis L, Vranes M, Kämper J, Schipper K, Göhre V**. **2016**. Genetic Manipulation of the Plant Pathogen Ustilago maydis to Study Fungal Biology and Plant Microbe Interactions. *Journal of visualized experiments: JoVE*: 1–9.

**Brachmann A, König J, Julius C, Feldbrugge M**. **2004**. A reverse genetic approach for generating gene replacement mutants in Ustilago maydis. *Molecular Genetics and Genomics* **272**: 216–226.

**Casimiro-Soriguer CS, Muñoz-Mérida A, Pérez-Pulido AJ**. **2017**. Sma3s: A universal tool for easy functional annotation of proteomes and transcriptomes. *Proteomics* **17**: 1–4.

**de Castro E, Sigrist CJA, Gattiker A, Bulliard V, Langendijk-Genevaux PS, Gasteiger E, Bairoch A, Hulo N**. **2006**. ScanProsite: detection of PROSITE signature matches and ProRule-associated functional and structural residues in proteins. *Nucleic acids research* **34**: W362–5.

**Eisenhaber B, Schneider G, Wildpaner M, Eisenhaber F**. **2004**. A sensitive predictor for potential GPI lipid modification sites in fungal protein sequences and its application to genome-wide studies for Aspergillus nidulans, Candida albicans, Neurospora crassa, Saccharomyces cerevisiae and Schizosaccharomyces pombe. *Journal of molecular biology* **337**: 243–253.

**Gasteiger E, Gattiker A, Hoogland C, Ivanyi I, Appel RD, Bairoch A**. **2003**. ExPASy: The proteomics server for in-depth protein knowledge and analysis. *Nucleic acids research* **31**: 3784–3788.

**Gattiker A, Gasteiger E, Bairoch A**. **2002**. ScanProsite: a reference implementation of a PROSITE scanning tool. *Applied bioinformatics* **1**: 107–108.

**Gillissen B, Bergemann J, Sandmann C, Schroeer B, Bölker M, Kahmann R**. **1992**. A two-component regulatory system for self/non-self recognition in Ustilago maydis. *Cell* **68**: 647–657.

**Hibbett DS, Binder M, Bischoff JF, Blackwell M, Cannon PF, Eriksson OE, Huhndorf S, James T, Kirk PM, Lücking R, *et al.*** **2007**. A higher-level phylogenetic classification of the Fungi. *Mycological research* **111**: 509–547.

**J S, E F, T M**. **1989**. Molecular Cloning: A Laboratory Manual (2nd ed.). *Cold Spring Harbor, NY Cold Spring Harbor Laboratory Press.*: 1–34.

**Jones DT, Taylor WR, Thornton JM**. **1992**. The rapid generation of mutation data matrices from protein sequences. *Computer applications in the biosciences : CABIOS* **8**: 275–282.

**Julenius K**. **2007**. NetCGlyc 1.0: prediction of mammalian C-mannosylation sites. *Glycobiology* **17**: 868–876.

**Krogh A, Larsson B, Heijne von G, Sonnhammer EL**. **2001**. Predicting transmembrane protein topology with a hidden Markov model: application to complete genomes. *Journal of molecular biology* **305**: 567–580.

**Lanver D, Müller AN, Happel P, Schweizer G, Haas FB, Franitza M, Pellegrin C, Reissmann S, Altmüller J, Rensing SA, *et al.*** **2018**. The Biotrophic Development of Ustilago maydisStudied by RNA-Seq Analysis. *The Plant Cell* **30**: 300–323.

**Loubradou G, Brachmann A, Feldbrugge M, Kahmann R**. **2001**. A homologue of the transcriptional repressor Ssn6p antagonizes cAMP signalling in Ustilago maydis. *Molecular Microbiology* **40**: 719–730.

**Marín-Menguiano M, Moreno-Sánchez I, Barrales RR, Fernández-Álvarez A, Ibeas JI**. **2019**. N-glycosylation of the protein disulfide isomerase Pdi1 ensures full Ustilago maydis virulence. *PLoS Pathogens* **15**: e1007687.

**Moreno-Sánchez I, Pejenaute-Ochoa MD, Navarrete B, Barrales RR, Ibeas JI**. **2021**. Ustilago maydis Secreted Endo-Xylanases Are Involved in Fungal Filamentation and Proliferation on and Inside Plants. *Journal of fungi (Basel, Switzerland)* **7**: 1081.

**Sperschneider J, Dodds PN**. **2022**. EffectorP 3.0: Prediction of Apoplastic and Cytoplasmic Effectors in Fungi and Oomycetes. *Molecular plant-microbe interactions: MPMI* **35**: 146–156.

**Steentoft C, Vakhrushev SY, Joshi HJ, Kong Y, Vester-Christensen MB, Schjoldager KT-BG, Lavrsen K, Dabelsteen S, Pedersen NB, Marcos-Silva L, *et al.* 2013**. Precision mapping of the human O-GalNAc glycoproteome through SimpleCell technology. *The EMBO Journal* **32**: 1478–1488.

**Varadi M, Anyango S, Deshpande M, Nair S, Natassia C, Yordanova G, Yuan D, Stroe O, Wood G, Laydon A, *et al.* 2022**. AlphaFold Protein Structure Database: massively expanding the structural coverage of protein-sequence space with high-accuracy models. *Nucleic acids research* **50**: D439–D444.

**Zuo W, Ökmen B, Depotter JRL, Ebert MK, Redkar A, Misas Villamil J, Doehlemann G**. **2019**. Molecular Interactions Between Smut Fungi and Their Host Plants. *Annual Review of Phytopathology* **57**: 411–430.
